# Towards a causal role of Broca’s area in language: A TMS-EEG study on syntactic prediction

**DOI:** 10.1101/2021.04.14.439631

**Authors:** Matteo Maran, Ole Numssen, Gesa Hartwigsen, Angela D. Friederici, Emiliano Zaccarella

## Abstract

Categorical predictions have been proposed as the key mechanism supporting the fast pace of syntactic composition in human language. Accordingly, grammar-based expectations facilitate the analysis of incoming syntactic information—e.g., hearing the determiner “the” enhances the prediction of a noun—which is then checked against a single or few other word categories. Previous functional neuroimaging studies point towards Broca’s area in the left inferior frontal gyrus (IFG) as one fundamental cortical region involved in categorical prediction during on-line language processing. Causal evidence for this hypothesis is however still missing. In this study, we combined Electroencephalography (EEG) and Transcranial Magnetic Stimulation (TMS) to test whether Broca’s area is functionally relevant in predictive mechanisms for language. Specifically, we transiently perturbed Broca’s area during the categorical prediction phase in two-word constructions, while simultaneously measuring the Event-Related Potential (ERP) correlates of syntactic composition. We reasoned that if Broca’s area is involved in predictive mechanisms for syntax, disruptive TMS during the processing of the first word (determiner/pronoun) would mitigate the difference in ERP responses for predicted and unpredicted categories when composing basic phrases and sentences. Contrary to our hypothesis, perturbation of Broca’s area at the predictive stage did not affect the ERP correlates of basic composition. The correlation strength between the electrical field induced by TMS and the magnitude of the EEG response on the scalp further confirmed this pattern. We discuss the present results in light of an alternative account of the role of Broca’s area in syntactic composition, namely the bottom-up integration of words into constituents.

## 1. INTRODUCTION

The combination of words into larger units is a hallmark of the human language faculty. A compositional engine overcomes the size of the lexicon, making it possible to convey an infinite number of meanings from a limited set of words. Syntactic rules are at the basis of this process, binding words into hierarchically structured phrases and sentences according to grammatical categorical information (Berwick et al., 2013; Chomsky, 1995; Everaert et al., 2015; Friederici et al., 2017).

At the neural level, the analysis of grammatical category is prioritized over other linguistic information (Friederici, 2011), mirroring the central role of syntactic composition in language. This is reflected in the earliness of the Event-Related Potential (ERP) components elicited by syntactic categorical violations (e.g., \**the forget*^1^), such as the Early Left Anterior Negativity (ELAN, Friederici et al., 1993; Neville et al., 1991), the Early Syntactic Negativity (ESN, Hasting & Kotz, 2008) and the Syntactic Mismatch Negativity (sMMN, Hasting et al., 2007; Herrmann et al., 2009). The latencies of these components show that categorical analysis occurs in approximately 200-250 milliseconds (ms), preceding thematic and semantic relations (Friederici, 2011). Syntactic effects are also observed on early perceptual components when the presence of closed-class morphemes or orthographic cues facilitates category recognition (Dikker et al., 2009, 2010). This first step of syntactic analysis occurs in a highly automatic fashion, as the sMMN and ESN effects are also elicited in the presence of distracting conditions (Batterink & Neville, 2013; Hasting et al., 2007; Hasting & Kotz, 2008; Herrmann et al., 2009). Moreover, the ELAN is not influenced by task-specific strategies or the probability of violation occurrence (Hahne & Friederici, 2002; Hahne & Friederici, 1999).

The earliness of categorical analysis has been proposed to rely on syntactic predictive mechanisms (Dikker et al., 2009, 2010; Jakuszeit et al., 2013; Lau et al., 2006). According to this hypothesis, syntactic predictions restrict the grammatical information to be checked against a target category (e.g., *The* → prediction for a noun) and allow fast analysis of incoming words.

Computationally, this idea is reminiscent of left-corner parsing models (Abney & Johnson, 1991; Hale, 2014; Resnik, 1992), which perform incremental syntactic analysis by opening a phrase as soon as its leftmost element is encountered (e.g., *The* → opening of a determiner phrase and prediction for a noun). At the neural level, this hypothesis is grounded on the assumption that the brain minimizes the processing load of incoming input by top-down predictions, which are passed from higher to lower levels of the functional architecture (Friston, 2003; Friston & Kiebel, 2009; Rao & Ballard, 1999). When the current input does not correspond to the expected one, a mismatch signal (i.e., the prediction error) is generated and the internal model is updated (Den Ouden et al., 2012; Friston, 2005; Garrido et al., 2007). Accordingly, the earliness of the ELAN would then reflect an incremental parsing process in which syntactic information of incoming words is checked against a single predicted candidate category (e.g., noun) or its left-side modifiers (e.g., adjectives). Overall, the use of structural information driving categorical expectations converges on data showing that preceding context facilitates different stages of linguistic analysis, including orthographic or phonological processing, lexical access and semantic integration (see Kuperberg & Jaeger, 2016 and Pickering & Gambi, 2018 for two recent reviews).

Evidence for the existence of categorical predictions in language comes from behavioural, neurophysiological and hemodynamic data. At the behavioural level, predictions driven by syntactic structure have been shown to influence fixation times when reading sentences (Boston et al., 2008). A second eye-tracking study, in which the final word of a sentence was displayed in one of two positions on the screen according to its grammatical category, found anticipatory eye-movements towards the correct target position, suggesting that participants used syntactic structures to inform categorical predictions (Bonhage et al., 2015). Going to the most basic two-word level, recent data showed faster recognition of target categories (noun or verb) when they were coherent with the expectation triggered by the primes (determiners and pronouns), independently of whether the primes were presented above or below awareness threshold (Berkovitch & Dehaene, 2019). At the neurophysiological level, Lau and colleagues (2006) showed that the amplitude of the ELAN depends on the strength of categorical predictions induced by the previous context. The authors observed that an ungrammatical sentence continuation elicited a smaller ELAN than the control condition when an ellipsis configuration softened the expectancy for a noun. Converging evidence comes from electroencephalography (EEG) studies employing narratives, which showed that metrics reflecting grammar-based expectations predict the signal elicited by incoming words (Brennan & Hale, 2019; Hale et al., 2018). Finally, at the two-word level increased oscillatory synchronization was observed before pseudo-verbs preceded by pronouns (Segaert et al., 2018), possibly reflecting the expectation for syntactic composition to occur with an upcoming verb element (see Lewis et al., 2015, 2016 and Meyer, 2018 for reviews on the oscillatory dynamics of linguistic analysis and prediction).

At the neuroanatomical level, syntactic violations are known to engage the left perisylvian cortex (Friederici et al., 2003; Herrmann et al., 2012; Suzuki & Sakai, 2003; Vandenberghe et al., 2002). Activity in these regions seems to be modulated by surprisal, a metrics reflecting how much the current grammatical information is unexpected given the previous context (Brennan et al., 2016; Henderson et al., 2016; Shain et al., 2020). However, it is unclear whether these studies isolated brain regions involved in generating or checking grammatical predictions. To the best of our knowledge, we are aware of only one study that directly investigated the generation of categorical prediction at neural level (Bonhage et al., 2015). In this experiment, the fMRI analysis was constrained by the timing of prediction generation, indicated by the anticipatory eye-movements towards the position of a target category. When only the structural information could be extracted from the context, increased activation as a function of syntactic prediction was observed in Broca’s area. Broca’s area is a region situated in the left inferior frontal gyrus (IFG), and its structural (Finkl et al., 2020) and functional (Trettenbrein et al., 2020) profile point towards a role in modality-independent linguistic computations, based on grammar (Chen et al., 2019; Chen, Goucha, et al., 2021; Chen, Wu, et al., 2021). This region is well-known to support linguistic composition, as shown by numerous fMRI studies (Graessner et al., 2021; Makuuchi et al., 2009; Pallier et al., 2011; Schell et al., 2017; Snijders et al., 2009; Tyler et al., 2010; van der Burght et al., 2019; Zaccarella, Meyer, et al., 2017), lesion data (Friederici et al., 1998, 1999) and meta-analytical findings (Hagoort & Indefrey, 2014; Zaccarella, Schell, et al., 2017). Within Broca’s area, the pars opercularis (Brodmann area, BA, 44) has been specifically linked to syntactic composition based on abstract categorical representations, as structure-building effects in this region are also observed during the processing of jabberwocky phrases or sentences (Goucha & Friederici, 2015; Zaccarella & Friederici, 2015a). Given that in jabberwocky conditions content elements (e.g., nouns, verbs, adjectives) are replaced by pseudowords, activity in BA44 might be amplified by the highly predictive nature of the functional elements (determiners, prepositions, morphological particles) retained within the stimuli. Furthermore, increased directed connectivity from BA44 to the left middle temporal gyrus (MTG) is observed when two-word phrases start with a function word compared to a non-predictive element (Wu et al., 2019), possibly reflecting the top-down transmission of a categorical expectation.

The hypothesis that Broca’s area and specifically BA44 is involved in generating categorical predictions appears to be coherent with computational parsing models and functional data from the neuroimaging literature, and is in line with the existence of domain specific circuits supporting predictive processes in language (Shain et al., 2020). However, conflicting evidence and theoretical views not supporting this notion have also being reported. First, words whose grammatical category is not expected, but which can nonetheless be integrated in a grammatical construction, do not seem to elicit an ELAN (Friederici et al., 1996). This early independence between grammaticality and predictive mechanisms has been also reported in sMMN-based studies, where the neural response to different grammatically correct phrases is not modulated by the frequency of occurrence of the phrase under analysis (Herrmann et al., 2009; Pulvermüller & Assadollahi, 2007). Secondly, given that increased activity in Broca’s area is also observed for syntactic categorical violations (Friederici et al., 2003; Herrmann et al., 2012), it is not possible to exclude that this brain region licenses syntactic structures via a bottom-up process rather than predict them. As a matter of fact, a very recent fMRI study showed that Broca’s area activity correlates with indexes of bottom-up integration during naturalistic listening (Bhattasali et al., 2019). Similarly, increased activity in the left IFG has been reported as a function of whether a word can be integrated or not in the syntactic context (Hultén et al., 2019). Third, recent data suggest that a careful examination of apparent pre- activation effects is necessary. While initial data supported the notion that probabilistic information can be used to anticipate properties of upcoming words up to the phonological level (Delong et al., 2005), a recent large-scale replication study showed that this effect might be absent or much smaller than originally thought (Nieuwland et al., 2018). Finally, theoretical views have emerged which put forward the notion that this process might not be a necessary component of linguistic comprehension (Huettig, 2015; Huettig & Mani, 2016).

At present, no causal evidence exists for or against the existence of categorical predictive processes located in Broca’s area. The absence of the ELAN in patients with lesions in Broca’s area (Friederici et al., 1998, 1999) supports a causal role of this region in syntactic composition, but does not discriminate between predictive and bottom-up processes. Both accounts are compatible with the absence of the ELAN, either because no categorical expectation is formed or because the integration phase is disrupted. A similar argument applies to a clinical study of Jakuszeit and colleagues (2013). Here we begin to address the computational role of Broca’s area in syntactic composition by testing one of the two competing hypotheses. In particular, we tested the causal role of Broca’s area in generating categorical predictions by using focal perturbations induced by short trains of Transcranial Magnetic Stimulation (TMS). When delivered “online” (i.e., during the task), TMS allows to test causal relationships between the targeted area and a specific cognitive process of interest (Hartwigsen, 2015; Pascual-Leone et al., 1999; Siebner et al., 2009; Walsh & Cowey, 2000). Our experiment represents the first investigation of the causal involvement of Broca’s area in generating syntactic predictions by combining three elements:

1. An ESN paradigm in which syntactic categorical predictions are generated at the basic two- word level (determiner → prediction for a noun, pronoun → prediction for a verb), and fulfilled (grammatical constructions) or violated (ungrammatical constructions).
2. An ERP analysis measuring the different brain responses to prediction fulfilment and violation.
3. A TMS approach with high temporal resolution to causally link Broca’s area to a specific stage of syntactic analysis (i.e., the generation of predictions).

We reasoned that, if Broca’s area is causally involved in syntactic predictive processes, TMS- induced disruption of this region during the prediction phase (determiner or pronoun processing) will attenuate the difference between expected and unexpected categories (nouns or verbs), which appear as second words in grammatical and ungrammatical constructions, respectively. More specifically we expect to find evidence for a Grammaticality*TMS interaction on the ESN amplitude, which we will further quantify using Bayesian statistics. At the most fine-grained level, we characterize the effect of TMS on categorical predictive processes by looking at the correlation between the strength of the electrical field induced by the stimulation in Broca’s area and changes in the ESN amplitude. If, conversely, our experiment will not support a causal role of Broca’s area in syntactic composition at the predictive stage, a reversed account relying on bottom-up integration processes can be instead proposed for the region.

## 2. METHODS

### 2.1 Participants

Thirty native German speakers were recruited for the experiment. Due to the presence of strong artifacts in the EEG signal, one subject was excluded from the analysis. Therefore, twenty-nine subjects were included in the statistical analysis (fifteen female; mean age: 27.1 years, standard deviation: 4.1 years). All participants were right-handed (mean laterality quotient: 93.3, standard deviation: 9.5), as assessed with the Edinburgh handedness test (Oldfield, 1971), had normal or corrected-to-normal vision, and no colour blindness. None of the participants presented contraindications against TMS or had history of psychiatric or neurological disorders. Participants gave their written informed consent and were reimbursed 12€ per hour for participating in the study. The study was approved by the local ethics committee (University of Leipzig) and was conducted in compliance with the Declaration of Helsinki guidelines.

### 2.2 Paradigm

Our experiment employed an adapted version of a standard two-word auditory ESN paradigm with syntactic categorical violations, previously published in the literature and frequently used to test neural sensitivity to syntactic composition (Hasting et al., 2007; Hasting & Kotz, 2008; Herrmann et al., 2009, 2012; Jakuszeit et al., 2013). The first word of each utterance was the German determiner “Ein” (*a*) or the personal pronoun “Er” (*he*), while the second word could be either a noun or verb. Thirty-two pairs of nouns and verbs with an ambiguous stem were used (e.g., “Fech- ter”, *fencer*, and “fech-tet”, *fences,* see section 2.3). Each second word was presented once following the determiner and once following the personal pronoun, resulting in four possible types of trials, two grammatical (*a* + *noun*, *he + verb*) and two ungrammatical (*a* + *verb*, *he + noun*). The grammatical and ungrammatical conditions consisted of sixty-four trials each, as thirty-two pairs of nouns and verbs were used. The conditions of the paradigm are summarised in Table 1.

**Table 1:** Conditions included in the experimental paradigm. The design crossed Grammaticality (grammatical vs. ungrammatical) and TMS (Brodmann Area (BA) 44, superior parietal lobe (SPL) and sham).

| Experimental conditions |  |
| --- | --- |
| Grammatical | Ungrammatical |
| Ein Falter (*a butterfly*) | Ein faltet (*a folds*) |
| Er faltet (*he folds*) | Er Falter (*he butterfly*) |

Grammaticality constitutes the first factor in our experimental design, reflecting whether the second word matched the categorical prediction triggered by the first one (grammatical items) or not (ungrammatical items). Importantly, with this paradigm, grammaticality is orthogonal to both the identity of the first word (“Ein” or “Er”) and the grammatical category of the second word (noun or verb), therefore ruling out potential confounding factors. As previously shown (Hasting & Kotz, 2008), ungrammatical items result in an increased ESN response, functionally equivalent to the

ELAN observed with longer stimuli (Friederici, 2011).

### 2.3 Stimuli

As in the original version of the paradigm (Hasting & Kotz, 2008), we used nouns and verbs with ambiguous stems in which the category information can only be assessed once the suffix is processed (e.g., “Fech-ter”, *fencer*, and “fech-tet”, *fences*) as second words. In this way, we could precisely time-lock the ERP analysis to the point of categorical access of the second word, which is represented by the suffix onset time. While in the original study (Hasting & Kotz, 2008) the category of most of the nouns was expressed with zero marking (e.g., “Kegel-Ø^2^”, *cone*, compared to the verb “kegel-t”, *bowls*), we decided to include only nouns with the category overtly marked by a suffix. Our decision was motivated by studies which showed no syntactic categorical violation effects when nouns with zero marking were used (Dikker et al., 2009; Herrmann et al., 2009).

Furthermore, syntactic violations realised with an offending suffix (e.g., “*Er kegel-st”, \**he bowl*) are more robust against conditions of reduced statistical power such as small or heterogeneous sample sizes than unmarked ones (e.g., “*Er Kegel- Ø”, \**he cone*, Jakuszeit et al., 2013).

The thirty-two pairs of nouns and verbs with ambiguous stems used in our experiment are the result of a four-step selection procedure. First, we extracted from CELEX corpus (Baayen, Piepenbrock, & Gulikers,1995) masculine and neuter German disyllabic nouns ending in “-er”. We used only masculine and neuter nouns in the nominative case as they follow the determiner “Ein”, while feminine nouns and other cases would have required explicit agreement processes. Secondly, for each noun we constructed a potential verb “candidate” in the infinitive form, by replacing the suffix “-er” with the infinite form ending “-en” (e.g., “Fecht-er”*, fencer* → “fecht-en”*, to fence*). For nouns ending with “-ler”, we constructed an additional infinite candidate form using the ending “- eln” (e.g., “Schwin-dler”, *cheater* → “schwin-deln”, *to cheat*). Nouns for which the respective verb candidate was not found in CELEX corpus were excluded at this step. In the third step, the verbs were inflected in the present tense third-person singular form. Pairs in which the verb became monosyllabic when inflected (e.g., “spinnen”, *to spin* → “spinnt”, *spins*) were excluded from the list. Finally, as in German “Ein” and “Er” can form compounds prefixing both nouns and verbs, we removed pairs of nouns and verbs in which the ungrammatical forms could exist as a compound (i.e., “ein+verb” or “er+noun”) according to the majority of eight independent native German speakers. The auditory stimuli used in the experiment were prepared adapting the cross-splicing procedure described by Hasting and Kotz (2008). For each pair of nouns and verbs a trained German native speaker was asked to read several times three utterances:

a. The correct determiner phrase (e.g., “Ein Fech-ter”, *A fencer*);
b. The correct verb phrase (e.g., “Er fech-tet”*, he fences*);
c. The stem embedded in a meaningless pseudo-word phrase (e.g., “Lub fech-tek”).

The recordings were acquired in a soundproof cabin using Audacity software (sampling rate: 44100 Hz). The most similar determiner phrase, verb phrase and pseudo-word phrase were then selected for the cross-splicing procedure. From the determiner phrase, the word “Ein” and the noun suffix (e.g., “-ter”) were extracted. The pronoun “Er” and the verb suffix (e.g,, “-tet”) were then extracted from the verb phrase. When the two first words (“Ein” and “Er”) were extracted from the recordings, also the silence extending up to 600 ms (the closest zero-crossing sample) from word onset was included. Similarly, the stem (e.g., “Fech”) was extracted from the pseudo-word phrase. To avoid clicking sounds, the recordings were cut only at points of zero crossing. The grammatical and ungrammatical utterances (e.g., |Ein |Fech|-ter|, *|Ein |Fech|-tet|, |Er |Fech|-tet|, *|Er |Fech|-ter|) were then created by concatenating one of the two first words (e.g., |Ein| or |Er|), the stem (e.g., |Fech|), and one of the two possible suffixes (e.g., |-ter|, |-tet|). Finally, the constructed utterances were normalized to 65 dB and 7 ms of silence were added at the beginning of each stimulus. Since TMS pulses produce a loud click noise, concatenated items were normalized to adjust the volume of the stimuli at the beginning of the experiment so that all the utterances could be heard clearly. Manipulation of the recordings was performed using Praat software (Boersma, 2001). Our procedure strongly reduced acoustic differences between grammatical and ungrammatical utterances up to the divergence point (DVP), after which the suffix occurs and the category of the second word is revealed (e.g., Ein Fech_[DVP]_ter, *Ein fech_[DVP]_tet, Er fech_[DVP]_tet, *Er Fech_[DVP]_ter). A t-test on the root mean square amplitude of the recordings up to the DVP revealed no significant difference between grammatical (Ein + Noun, Er + Verb) and ungrammatical (Ein + Verb, Er + Noun) items (*p =* .99).

### 2.4 Transcranial Magnetic Stimulation (TMS)

Transcranial magnetic stimulation was applied during the task (“online”) to investigate the causal role of BA44 in syntactic predictive processes. We delivered 10 Hz trains of five TMS pulses during the first word of each item (“Ein”, *a*, or “Er”, *he*) to perturb the stage of syntactic categorical prediction (determiner → prediction for a noun, pronoun → prediction for a verb). The first pulse of each TMS train was time-locked to the onset of the first word and each burst lasted 400 ms. Since the second word of each item started 600 ms after the first word onset and potential after-effects of online TMS are thought to last approximately half of the stimulation time (Rotenberg, Horvath, & Pascual-Leone, 2014), our stimulation protocol and stimuli materials ensured that the perturbation was limited to the stage of syntactic prediction only.

We included three TMS conditions: BA44 (MNI: -48, 17, 16), the left superior parietal lobe (SPL, MNI: -34, -42, 70) as an active control site, and a sham condition (Figure 1). Each participant took part in three experimental sessions, one for each TMS condition, which were separated at least by 7 days (mean distance: 7.89 days, standard deviation: 2.96 days). The order of conditions was counterbalanced across subjects. The MNI coordinates for BA44 were defined according to the results of Zaccarella and Friederici (2015), who found increased activation for phrases compared to word lists in this region. This target is located in the most anterior and ventral part of BA44, which is functionally specialized in syntactic computations (Papitto et al., 2020; Zaccarella et al., 2021). The SPL coordinates were based on a TMS experiment on degraded speech comprehension in which this region served as a control condition (Hartwigsen et al., 2015). In the sham condition, no effective stimulation of the brain occurred. The vertex was chosen as a target for sham TMS to perform the neuronavigation procedure as in the other TMS conditions (Friehs et al., 2020; Klaus & Hartwigsen, 2019; Kuhnke et al., 2020). In the sham condition, a disconnected coil was navigated over the electrode Cz and an active coil was placed above it with an angle of 90°, therefore not stimulating the brain. This procedure allows to produce the same acoustic noise as the other two TMS conditions, but without an actual stimulation of the brain (Friehs et al., 2020; Kroczek et al., 2019; Kuhnke et al., 2017, 2020; Meyer et al., 2018).

**Figure 1:**
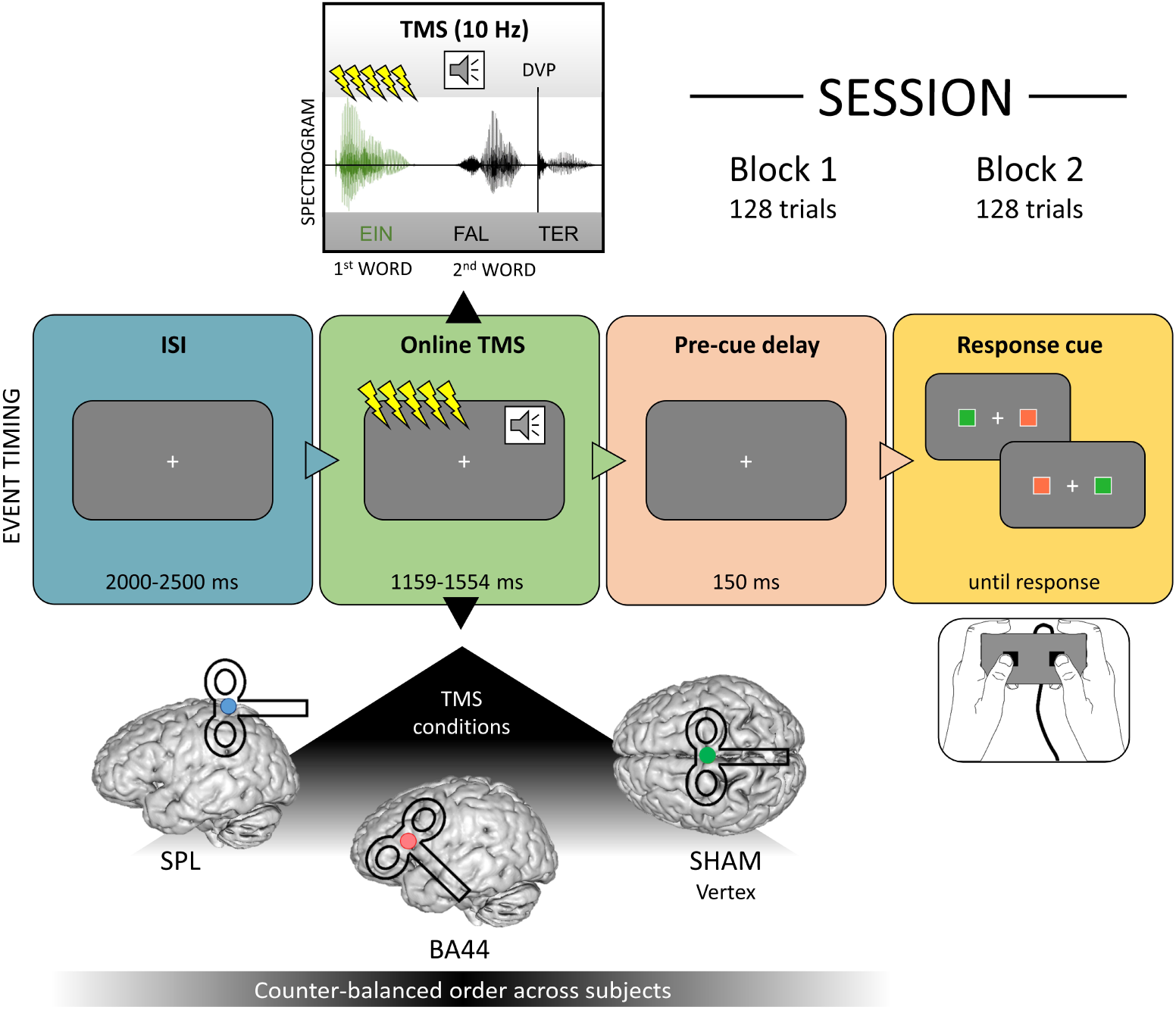
Timing of events including an illustration of the online stimulation during the first word (above the event timeline) and the three target regions for the neuro-navigation system (below the event timeline). Divergence Point (DVP); Interstimulus Interval (ISI); Superior parietal lobe (SPL); Brodmann Area (BA) 44.

We used stereotactic neuronavigation (TMS Navigator software version 3.0.33, Localite GmbH, Sankt Augustin, Germany) to position and maintain the TMS coil over the target regions during the experiment. Individual structural T1-weighted MRI images were previously acquired for each participant. The coordinates of the target regions were converted from the MNI standard to the individual subject space using SPM12 software (Wellcome TrustCenter for Neuroimaging, University College London, United Kingdom), using an established procedure (Friehs et al., 2020; Klaus & Hartwigsen, 2019; Kuhnke et al., 2017, 2020). After EEG preparation, the head of each participant was co-registered to their MRI image, allowing for precise positioning of the TMS coil over the target coordinates as defined in the individual anatomical image.

TMS was delivered using a figure-of-eight coil (C-B60) connected to a MagVenture MagPro X100 stimulator (MagVenture, Farum, Denmark). The coil handle was oriented with an angle of 45° and 0° relative to the sagittal plane when stimulating BA44 and the SPL respectively, as in previous studies (Hartwigsen et al., 2015; Klaus & Hartwigsen, 2019; Kroczek et al., 2019; Kuhnke et al., 2017; Meyer et al., 2018). The intensity of the stimulation was set to 90% of the individual resting motor threshold (RMT), defined during the first session for each participant. RMT was defined as the minimum intensity at which TMS could evoke at least 5 motor evoked potentials (MEP) with an amplitude ≥ 50 µV in the relaxed first dorsal interosseous muscle out of 10 consecutive pulses (Rothwell et al., 1999). To this end, the TMS coil was navigated over the coordinates of the left hand motor area (MNI: –37, –21, 58, Mayka et al., 2006) and the hotspot was identified with a standard threshold hunting procedure. If necessary, stimulation intensity for BA44 was corrected for the scalp-to-cortex distance relative to the motor cortex as described previously (e.g., Kuhnke et al., 2017). In short, the adjusted intensity was calculated using the formula by Stokes and colleagues (2005), as adapted for applications with 90% of the RMT (Kuhnke et al., 2017): *BA44 intensity (stimulator output) = 90% RMT+3*(Distance*_BA44_ *– Distance*_M1_), where Distance_BA44_ and Distance_M1_ correspond to the distance in mm between the scalp and BA44 and M1 respectively. The stimulation intensity for the sham condition was the same as the one used for BA44. Finally, the stimulation intensity for the SPL condition corresponded to the 90% of the RMT, as for no subject it required to be adjusted. If stimulation was too unpleasant, intensity was gradually decreased in steps of 1%.

### 2.5 Procedure and timing of events

At the beginning of the TMS-EEG sessions, each participant filled in a short TMS safety questionnaire and received the task instructions After EEG preparation, the participant was moved into an electrically shielded cabin where the head surface was co-registered to the structural MRI image for TMS neuronavigation. Subjects sat comfortably approximately 140 cm from the computer monitor. During the first TMS-EEG session, the individual RMT was defined. To familiarize the subject with the sensory stimulation associated with each TMS condition, some test pulses on the target region were delivered. Before the experiment, subjects were provided with in- ear headphones and after reading a reminder of the instructions they underwent a short practice, consisting of 12 trials with items excluded from the main task. The trial structure of the practice block was the same as the one of the main task, but feedback was provided after each response to ensure that subjects understood the instructions. To provide comparable conditions, TMS was also delivered during the practice trials, allowing the subject to indicate if sound volume needed to be adjusted due to the TMS-induced noise.

During the task, the TMS coil was manually positioned and maintained over the target region. Subjects performed a grammaticality judgement task, indicating if the two-word utterance they heard was grammatically correct or not via a button-box press. A fixation cross was displayed at the centre of the monitor, and after an inter-stimulus interval randomly jittered between 2 and 2.5 seconds (s) the two-word item was presented acoustically. The TMS train was delivered during the first word. After the acoustic item ended, a delay of 150 ms was included to avoid an overlap of language-related and motor-related evoked responses in the EEG signal. A response cue was then presented, consisting of two coloured squares presented to the left and right of the fixation cross. One of the squares was red and one green, with the colours being assigned pseudo-randomly for each trial. The green colour was associated to the position of the response button for “grammatical”, similarly the red colour coded for “ungrammatical”. We used a red and a green colour with a similar luminance (L = 64.39 and 64.37 respectively in CieLuv color-space) to avoid that differences in brightness might bias the behavioural data analysis. Relative luminance was calculated implementing the formula defined in the Web Content Accessibility Guidelines (WCAG) 2.0 (https://www.w3.org/TR/WCAG20/Overview.html#sRGB, see Supplementary Materials). The timing of TMS bursts and stimulus presentation was controlled using Presentation Software version 17.2 (NeurobehavioralSystems, Inc., Albany, CA, USA). Figure 1 illustrates the structure of a trial. In each session, subjects performed the task twice (in two blocks), with the same 128 items presented in a different pseudo-randomized order. Short breaks were included every 32 trials, to cool down and switch the TMS coil if needed. After the experiments, the position of the electrodes was digitized using the TMS Navigator software. Considering EEG-TMS preparation, the first experimental session lasted on average approximately 3.5 hours, while the other two lasted approximately 2.5 hours.

### 2.6 Behavioural data analysis

Behavioural data were analysed with repeated measures analysis of variance (ANOVA), including as factors Grammaticality (grammatical and ungrammatical), TMS (BA44, SPL and sham) and Block (first and second). Repeated measures ANOVA was conducted on the subjects’ mean responses times and accuracy rates for each condition, following the removal or trials with RTs shorter than 150 ms or longer than 1 s. Analysis of the RTs was based on trials with correct response only.

#### 2.7.1 EEG recording and analysis

TMS pulses result in a series of artifacts on the concurrent EEG signal which need to be controlled during both data collection and pre-processing. Electromagnetic artifacts are commonly observed following each TMS pulse (Ilmoniemi & Kičić, 2010; Rogasch et al., 2013, 2014; Veniero et al., 2009), and depending on the target location additional large cranial muscular activity can contaminate the EEG signal. Muscle artifacts are particularly pronounced when the target site is a lateral brain region (Mutanen et al., 2013; Rogasch et al., 2013), such as the IFG and the posterior temporal lobe (Salo et al., 2020). Given the series of potential artifacts during online TMS-EEG, the employed procedure for data collection and pre-processing in the present study differs from traditional EEG studies.

EEG signal was recorded using 63 Ag/AgCl monopolar electrodes (61 electrodes embedded in an EEG cap, EC80, EasyCap GmbH, Germany, and A1 and A2 on the left and right mastoids respectively), which were placed according to the international extended 10-20 system. Two additional pairs of bipolar electrodes were placed to monitor vertical and horizontal eye movements. EEG signal was amplified using REFA8 68-channel amplifier system (TMSi, Oldenzaal, the Netherlands) and recorded at a sampling rate of 2000 Hz using BrainVision Recorder software version 1.02.0001 (Brain Products GmbH, Gilching, Germany). The average of the 63 monopolar electrodes served as an online reference. The ground electrode was placed on the sternum. Electromagnetic artifacts following each TMS pulse were reduced by arranging the direction of the electrode wires orthogonally to the TMS coil handle (Sekiguchi et al., 2011). Impedance was kept below 5 kΩ.

Pre-processing was performed using the Matlab FieldTrip toolbox version fieldtrip-20200115 (Oostenveld et al., 2011). Given that the TMS trains were time-locked to the first word onset, EEG signal in this time-window was strongly contaminated by the large electromagnetic and muscular artifacts described at the beginning of this section. The presence of these artifacts could have resulted in large signal distortions when applying common EEG pre-processing steps like filtering on the raw data (Rogasch et al., 2017). Since our ERP component of interest is time-locked to the DVP of the second word, we applied cubic interpolation of the continuous EEG signal from -2 to 450 ms relative to the first pulse of each TMS train (first word onset). Cubic interpolation was based on the 300 ms time-window before and after the segments to be interpolated. The continuous EEG signal obtained after interpolation was high-pass filtered with a cutoff frequency of 0.5 Hz (onepass-zerophase, order 4460, kaiser-windowed sinc FIR, 6 dB attenuation at the cutoff frequency, transition width 1.0 Hz, stopband 0-0.0 Hz, passband 1.0-1000 Hz, max passband deviation 0.0100, stopband attenuation 40 dB). Epochs from -250 ms to 2 s relative to the DVP of the second word were then extracted. Participants were instructed to delay, if possible, blinks and eye-movements until after the behavioural response, however ocular artifacts were present also in earlier parts of the trial. Therefore, this extended time-window for epoching allowed us to have a sufficient number of blinks and eye-movement events for well characterizing ocular artifacts with Independent Component Analysis (ICA) in a later step. Before ICA, epochs were visually inspected and trials and channels with excessive artifacts were removed (trials removed per block: mean = 3.4, std = 3.5; channels removed per block: mean = 0.9, std = 0.9). The common average reference of the good channels was then computed, and ICA using the RunICA algorithm was run, accounting for data rank reduction due to bad channel exclusion. ICA components were visually inspected and bad components reflecting ocular, cardiac and muscle artifacts were removed. If present, components reflecting the exponential decay after TMS were removed as well. After the removal of bad ICA components (number of components kept per block: mean = 27.1, standard deviation = 5.2), EEG data were re-referenced to the common average and the signal of the channels removed during visual inspection was interpolated using spherical spline interpolation (Perrin et al., 1989), an approach recently used in a TMS-EEG experiment targeting Broca’s area (Kroczek et al., 2019). EEG data were then re-referenced to the new common average reference and trials with an incorrect response were removed. The clean trials with a correct response were low-pass filtered with a cut- off frequency of 44 Hz (onepass-zerophase, order 408, kaiser-windowed sinc FIR, 6 dB attenuation at the cutoff frequency, transition width 11.0 Hz, passband 0-38.5 Hz, stopband 49.5-1000 Hz, max passband deviation 0.0100, stopband attenuation 40 dB). These prprocessing steps were repeated for each of the two blocks in each session. The trials from the two blocks were then merged in one unique dataset per TMS condition for each subject and re-referenced to the average of A1 and A2 electrodes. No baseline correction was applied, as the use of our high-pass filter already attenuated direct-current offset (Widmann et al., 2015). From each dataset two ERP waveforms were then calculated, averaging separately the trials belonging to the grammatical and ungrammatical conditions. This procedure resulted in six ERP waveforms per subject, reflecting the six cells of our Grammaticality*TMS within-subject design. ERP waveforms were then calculated to test three effects of interest: the main effect of Grammaticality (averaged across TMS conditions), the main effect of TMS (averaged across stimulus conditions in each session) and the interaction between Grammaticality and TMS.

The statistical analysis of EEG data was performed using non-parametric cluster-based permutation tests (Maris & Oostenveld, 2007) implemented in the FieldTrip toolbox (Oostenveld et al., 2011). This method is based on cluster formation according to test values and spatial and temporal contiguity after sample-by-sample statistical comparison. A cluster-level statistic is then calculated and, by comparing it against its distribution in random partitions via the Monte Carlo approximation, a significance probability value is obtained (for a detailed presentation, see Maris & Oostenveld, 2007). The dependent sample T-statistic (“depsamplesT”) was used for cluster formation when analysing the main effect of Grammaticality. For the analysis of the main effect of TMS and the Grammaticality*TMS interaction, the dependent sample F-statistic (“depsamplesFunivariate”) was used, as three levels were present in the independent variable^3^. The cluster-level statistic was calculated as the maximum of the cluster-level summed T- or F-values of each cluster. The critical alpha level for the Monte Carlo significance probability was set to 0.025 when testing the main effect of Grammaticality (two-tailed hypothesis) and to 0.05 for the analysis of the main effect of TMS and the Grammaticality*TMS interaction (one-tailed hypothesis). In each of the three statistical tests conducted (two main effects and one interaction), the Montecarlo estimation was based on 5000 random partitions and the time-window of interest was defined from 0 to 1000 ms relative to the DVP.

#### 2.7.2 Bayesian repeated measures ANOVA on the ESN amplitude

To quantify the evidence for and against the presence of a Grammaticality*TMS interaction in our EEG data, we performed an additional Bayesian repeated measures ANOVA on the mean amplitude of the ESN. Bayesian analysis allows to quantify evidence for both the null and the alternative hypotheses, describing how informative data from a given experiment are (Keysers et al., 2020; Wagenmakers, Marsman, et al., 2018). Bayes factors (BF) indicate how likely the data are under these two hypotheses. For example, a BF_10_ equal to 5 indicates that the current data are five times more likely under the alternative than the null hypothesis. BF_01_ is equal to 1/BF_10_ and indicates how many times the data are more likely under the null hypothesis.

In a Bayesian repeated measures ANOVA, Bayes Factors are obtained by comparing the predictive performance of two models (van den Bergh et al., 2020; Wagenmakers, Love, et al., 2018). Bayes Factors BF_10_ and BF_01_ quantify how much the data are more likely according to one of the two competing models (e.g., an alternative model against the null model or the best model). For example, a BF_10_ = 5 relative to a null model (e.g., a model that only accounts for the presence of different subjects) means that the data are predicted five times better by the given alternative model. A BF_01_ = 5 relative to a null model means that the null model predicts the data five times better than the alternative one.

The analysis was conducted using JASP software version 0.14 (JASP Team, 2020; https://jasp-stats.org/; for theoretical and practical introductions see Dablander et al., 2020; Faulkenberry et al., 2020; Keysers et al., 2020; Wagenmakers, Love, et al., 2018; Wagenmakers, Marsman, et al., 2018). The Bayesian repeated measures ANOVA included Grammaticality and TMS as factors. This analysis compared the performance of five models: a null model (M0: coding only the presence of different subjects) and four alternative models (M1: subject + Grammaticality, M2; subject + TMS, M3: subject + Grammaticality + TMS, M4: subject + Grammaticality + TMS + Grammaticality*TMS). The default uninformed prior distribution was used. We planned to test the Grammaticality*TMS interaction in two ways:

1. By comparing model M4 including the interaction against the models which included only the main effect of Grammaticality (M1) and the two main effects (M2). This comparison quantifies how much adding an interaction term improves the predictive performance of the model.
2. By performing an analysis of the effects via Bayesian Model Averaging, which allows to quantify the evidence for including factors and interactions by considering all the models according to their predictive power (Hinne et al., 2020; Keysers et al., 2020; van den Bergh et al., 2020; Wagenmakers, Love, et al., 2018). With this analysis BF_incl_ and BF_excl_ are obtained, indicating respectively how much more likely the data are under models which include and exclude a given factor or interaction. The analysis of effects was computed across all models.

For the Bayesian repeated measures ANOVA, we extracted the mean amplitude of the ESN component averaging signal between 190 ms and 430 ms at 40 electrodes: AF3, AFz, AF4, F5, F3, F1, Fz, F2, F4, F6, FC5, FC3, FC1, FCz, FC2, FC4, FC6, C5, C3, C1, Cz, C2, C4, C6, CP5, CP3, CP1, CPz, CP2, CP4, CP6, P5, P3, P1, Pz, P2, P4, P6, PO3, POz and PO4. The electrodes and time- points included are based on the results of the main effect of Grammaticality and by the rather spread topography of our ERP component of interest (see section 3.2 below). Henceforth we refer to this as the Full ESN. Crucially, the criterion used for selecting the electrodes and time-points included does not make circular the analysis, which addresses a different research question (interaction) compared the test used for defining them (main effect of Grammaticality).

#### 2.7.3 ESN and induced electrical field simulation

Together with stimulation intensity and coil orientation (Laakso et al., 2014; Weise et al., 2020), neuroanatomical factors such as individual gyrification patterns (Thielscher et al., 2011) and the distribution of tissue types (Lee et al., 2018; Opitz et al., 2011) affect the spread and strength of the electrical field induced by TMS pulses. To precisely characterize the impact of BA44 stimulation on the amplitude of the ESN, we performed an additional analysis on the EEG data including the strength of the electrical field in this target region for each subject. By modelling the extent to which TMS interfered with the target region it is possible to account for anatomical factors (Lee et al., 2018; Thielscher et al., 2011) which, differing between subjects, might otherwise hide the presence of an effect of TMS if not included in the analysis (Kuhnke et al., 2020).

The calculation of the induced electrical fields was implemented using a recently established pipeline (Weise et al., 2020). For each subject and each active TMS condition we performed an electrical field simulation based on individual T1-weighted images, additional T2-weighted images if available, and the coil position recorded during the experimental session. Individual head meshes were constructed using the headreco pipeline (Nielsen et al., 2018) and Simnibs software (Windhoff et al., 2013) was used to calculate the electric fields. The electric field models were visually inspected to ensure good quality of the head models. At this stage, two subjects were excluded from the analysis, due to an unrealistic field reconstruction. For each of the remaining 27 subject we extracted the average electrical field intensity from nine regions of interest (ROIs), two in Broca’s area (BA44 and BA45; Amunts et al., 1999, 2004), and seven in the SPL (BA5L, BA5M, BA5Ci, BA7A, BA7PC, BA7M, BA7P; Scheperjans, Eickhoff, et al., 2008; Scheperjans, Hermann, et al., 2008) using maximum probability maps from the SPM Anatomy Toolbox version 2.2c (Eickhoff et al., 2005, 2006, 2007). The inclusion of BA45 as a ROI is motivated by its involvement, together with BA44, in categorical prediction (Bonhage et al., 2015) and by its close proximity to this region in the left IFG^4^. The average electrical fields in Broca’s area (BA44 and BA45) and in SPL (BA5L, BA5M, BA5Ci, BA7A, BA7PC, BA7M, BA7P) ROIs were extracted from the BA44 and SPL TMS sessions respectively.

To test whether TMS affected the ESN, we computed a Pearson correlation between the induced electrical field in the abovementioned ROIs and the sham-normalized amplitude of Full ESN. The two sham-normalized Full ESN amplitudes were obtained in a two-step procedure:

1. First, for all the three TMS conditions we calculated the mean amplitude of the difference wave (ungrammatical – grammatical), resulting in three mean amplitude values: Full ESN_BA44_, Full ESN_SPL_ and Full ESN_sham_;
2. We then obtained the sham-normalized mean amplitudes (Full ESN_BA44_ effect, Full ESN_SPL_ effect) by subtracting Full ESN_sham_ from Full ESN_BA44_ and Full ESN_SPL_ respectively (Full ESN_BA44_ effect = Full ESN_BA44_ - Full ESN_sham_). As the induced electrical field for the sham condition is zero (no electrical stimulation of the brain), this subtraction isolated the effect of the induced field in a given ROI on the ESN amplitude for each of the two active TMS conditions.

Additionally, as our main effect of Grammaticality is characterized by an early frontal component and a second centro-parietal component (see Results section and Figure 3), we performed an exploratory analysis focusing on each component separately. This additional analysis is motivated by ERP studies showing the presence of two subsequent negativities for agreement (Barber & Carreiras, 2005; Hanna et al., 2014; Jakuszeit et al., 2013; Pulvermüller & Shtyrov, 2003) and categorical (Hasting et al., 2007) marked syntactic violations at the two-word level, which might reflect different stages of analysis. Crucially, by analysing the two effects separately we could test whether TMS selectively affected only one of them. We subdivided the Full ESN in two parts:

1. First ESN: average of signal from 190 ms to 310 ms at 17 anterior electrodes AF3, AFz, AF4, F5, F3, F1, Fz, F2, F4, F6, FC5, FC3, FC1, FCz, FC2, FC4 and FC6. The time-points included correspond to the first half of the Full ESN effect. Only anterior electrodes are included, in light of the topography of the main effect of grammaticality in this time- window (see Figure 3);
2. Second ESN: average of signal from 310 ms to 430 ms at 17 posterior electrodes CP5, CP3, CP1, CPz, CP2, CP4, CP6, P5, P3, P1, Pz, P2, P4, P6, PO3, POz and PO4 (second half of the time window of the Full ESN effect). Only posterior electrodes are included, since in this time-window the effect is mostly pronounced at these sites (see Figure 3).

**Figure 3:**
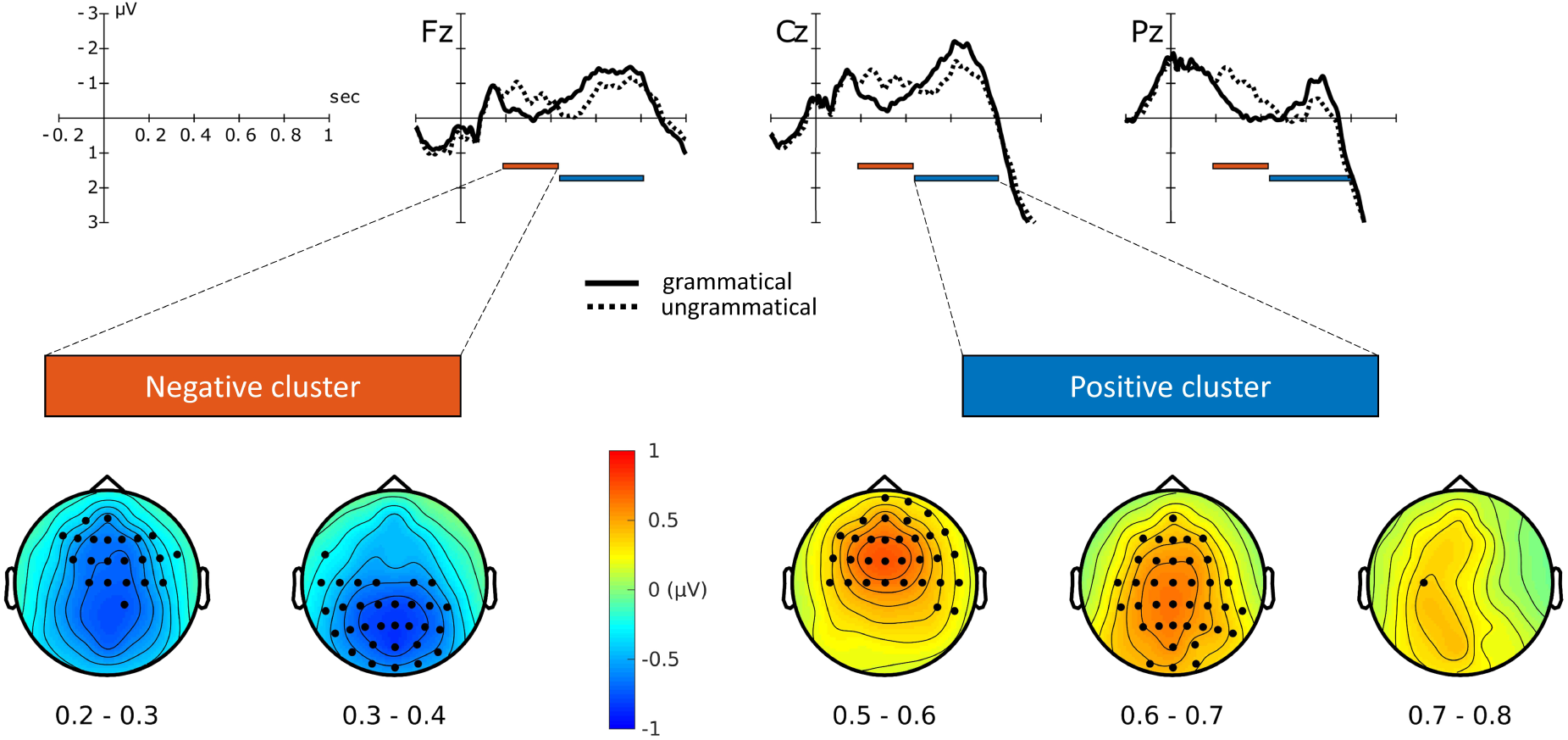
ERP waveforms for grammatical and ungrammatical waveforms (µV over seconds, collapsed across TMS conditions at selected electrodes), together with electrodes and time-points showing highest contributions to the significance of the negative (bottom left) and positive (bottom right) clusters.

First ESN_BA44_ effect, First ESN_SPL_, Second ESN_BA44_ effect and Second ESN_SPL_ effect were obtained with the same procedure described above for the full time-window, normalizing First ESN_BA44/SPL_ and Second ESN _BA44/SPL_ with the subtraction of First ESN_sham_ and Second ESN_sham_ respectively. The NHST correlational analysis was complemented by Bayesian inference using JASP software (JASP Team, 2020), to quantify both evidence for the alternative and the null hypotheses. The default uninformed prior distribution was used.

## 3. RESULTS

### 3.1 Behavioural data

The performance of the participants was at ceiling (mean accuracy = 97%, range = 75-100%), and the analysis of the accuracy revealed no significant main effect or interaction involving the factors Grammaticality, TMS and Block. The analysis of response times (RTs) showed a main effect of Grammaticality (*F*(1,28) = 92.43, *p* < 5e-10, η^2^*G* = 0.0655), with RTs for the grammatical items being on average 35 ms faster than for the ungrammatical ones. The main effect of Block was significant (*F*(1,28) = 11.35, *p* < 0.005, η^2^*G* = 0.0097), with RTs being on average 13 ms faster in the second block. Finally, the interaction Grammaticality*Block was significant (*F*(1,28) = 7.20, *p* < 0.05, η^2^*G* = 0.0008). A post-hoc analysis revealed that this interaction was driven by a significant difference between the RTs for the ungrammatical conditions of Block 1 and Block 2 (*p* < 0.001, Bonferroni-corrected), which was absent for the grammatical counterpart (*p* > .05, Bonferroni-corrected). No main effect of TMS and no interaction including this factor was significant. Figure 2 illustrates the results of the repeated measures ANOVA.

**Figure 2:**
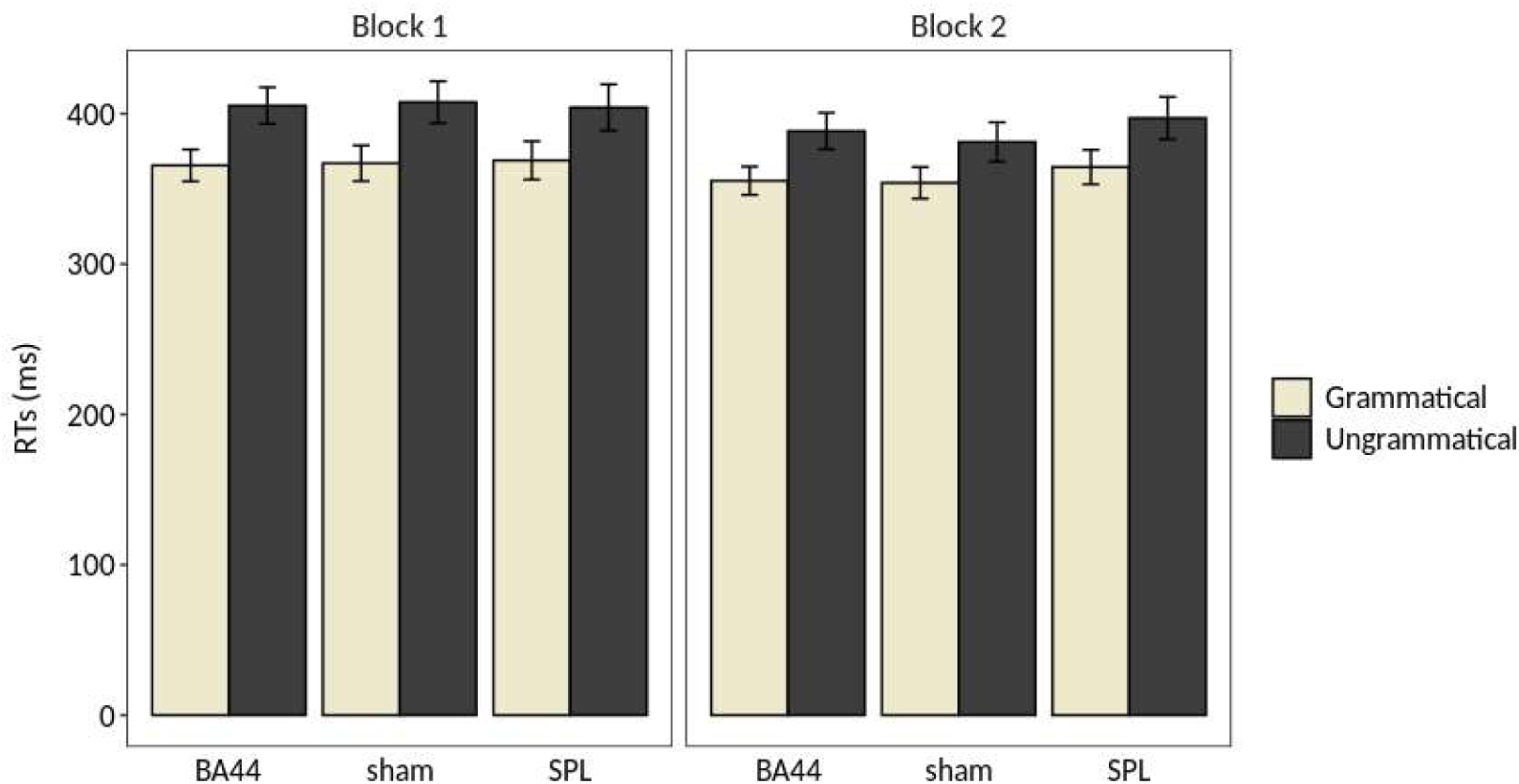
Results of the repeated measures ANOVA on the Response Times (RTs). The error bar indicates the standard error of the mean.

### 3.2 EEG data

The ERP waveforms of grammatical and ungrammatical conditions at selected electrodes, collapsed across TMS sites, are shown in Figure 3. Additional electrodes are displayed in the Supplementary Materials (Figure S1). Visual inspection of the ERP waveforms reveals an increased negativity for the ungrammatical condition from approximately 200 ms to 450 ms, followed by a late positivity from 450 to 800 ms. The cluster-based permutation test revealed a main effect of Grammaticality, with the presence of a significant negative (*P* < 0.0005, cluster-corrected) and positive (*P* < 0.0005, cluster-corrected) cluster. The negative cluster, reflecting increased negativity for the ungrammatical condition relative to the grammatical one, extended approximately from 190 to 430 ms after the DVP^5^ (Figure 3). The positive cluster, reflecting an effect in the opposite direction, extended approximately from 440 to 800 ms after the DVP. Both the effects were mostly pronounced over fronto-central and centro-parietal electrodes (Figure 3). A marginally non- significant effect of TMS was also found (*P* = 0.05, cluster-corrected).

The Grammaticality*TMS interaction of interest was not significant (*P >* 0.5, cluster-corrected). The ERP waveforms of grammatical and ungrammatical conditions within each TMS are shown in Figure 4. Additional electrodes are displayed in Figures S2, S3 and S4 in the Supplementary Materials. The absence of the interaction is evidenced by the presence of an increased negativity and a late positivity for the ungrammatical condition in each TMS condition. Indeed, within each TMS condition significant negative and positive clusters were found (BA44: first negative cluster *P* < 0.005, second negative cluster *P* < 0.005, first positive cluster *P* < 0.005, second positive cluster *P* < 0.05; sham: negative cluster *P* < 0.0005, positive cluster *P* < 0.0005; SPL: negative cluster *P* < 0.0005, positive cluster *P* < 0.005). The extent of the clusters in two selected time-windows is shown in Figure 4. The full extent of the clusters within each TMS condition is shown in Figures S5, S6, S7 and S8 in the Supplementary Materials. The absence of the critical Grammaticality*TMS interaction shows that TMS over Broca’s area during the first word did not affect the amplitude of the ESN.

**Figure 4:**
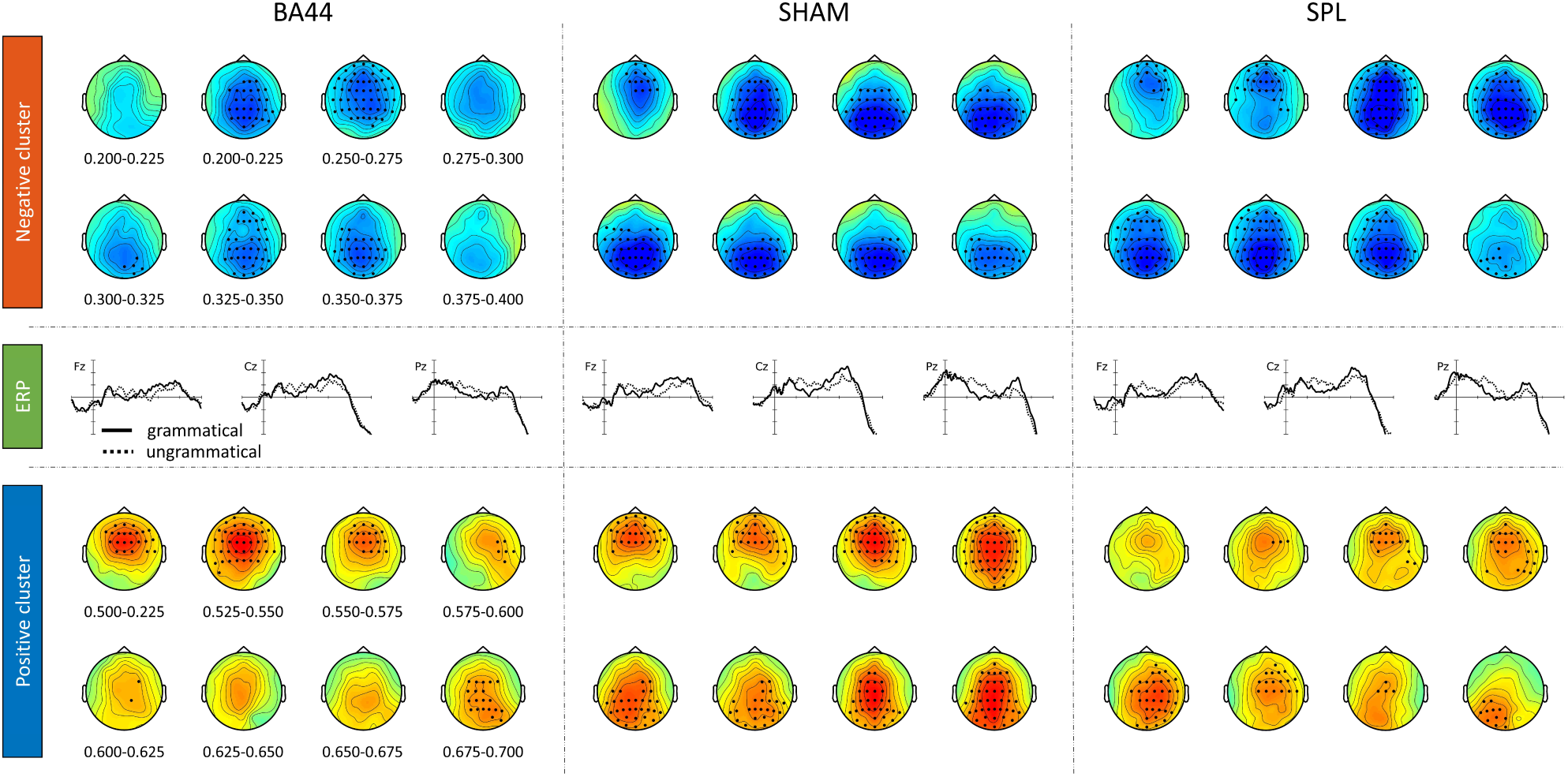
Grammaticality effect within each TMS condition. Electrodes and time-points providing the highest contribution to the significance of the negative cluster (top rows, 0.2-0.4 s time-window) and positive clusters (bottom rows, 0.5-0.7 s time-window) are highlighted. Colorbar and axes dimensions are the same as the ones shown in Figure 3.

Visual inspection of the ERP waveforms of the SPL condition shows an increased positivity for the ungrammatical items approximately 50 ms before the DVP. Crucially, this difference is short-lived, with the waveforms of grammatical and ungrammatical condition being aligned approximately 30 ms after the DVP. Furthermore, in the other two TMS conditions the waveforms are already aligned before the DVP. Differences between conditions can be problematic if they are sustained effects and are “masked” by baseline-correction procedure (Steinhauer & Drury, 2012). We did not perform a baseline-correction procedure, which would have artificially increased the Grammaticality effect for this TMS condition. To statistically confirm that this pre-DVP difference in the ERP waveforms is not a sustained effect, we performed a cluster-based permutation test on grammatical and ungrammatical conditions in the time-window -5 to 180 ms relative to the DVP. This analysis revealed no significant cluster (*P* > 0.5), in line with the non-sustained nature of this difference.

### 3.3 Bayesian repeated measures ANOVA on the ESN amplitude

The results of the Bayesian repeated measures ANOVA on the ESN amplitude are summarized in Table 2. The best model included only the factors subject and Grammaticality (BF_M_ = 5.651). The model including the Grammaticality*TMS interaction was approximately 10 times less likely than the model with only the main effect of Grammaticality given the data (BF_01_ = 10.295). Direct comparison of the interaction model against the one including the two main effects showed that the former was approximately 6 times less likely given the data (BF_01_ = 6.287).

**Table 2:** Summary of the results of the Bayesian repeated measure ANOVA conducted on the Full ESN. P(M) = prior model probability; P(M|data) = posterior model probability; BF_M_ = posterior model odds; BF_10_ and BF_01_ show the Bayes factors for the comparison of each model against the best one (Grammaticality).

| Models | P(M) | P(M\|data) | BF<sub>M</sub> | BF<sub>10</sub> | BF<sub>01</sub> | Error % |
| --- | --- | --- | --- | --- | --- | --- |
| Grammaticality (Gram) | 0.200 | 0.586 | 5.651 | 1.000 | 1.000 | - |
| Gram + TMS | 0.200 | 0.358 | 2.226 | 0.611 | 1.683 | 2.829 |
| Gram + TMS + Gram*TMS | 0.200 | 0.057 | 0.241 | 0.097 | 10.295 | 2.985 |
| Null model | 0.200 | 3.145e-9 | 1.258e-8 | 5.371e-9 | 1.862e+8 | 2.521 |
| TMS | 0.200 | 1.073e-9 | 4.291e-9 | 1.832e-9 | 5.459e+8 | 2.634 |

The analysis of the effects is summarized in Table 3. The data are approximately 1.5*10_7_ times more likely under models which include the Grammaticality factor (BF_incl_ = 1.539e+8) and two times more likely under models which do not include the TMS factor (BF_excl_ = 2.047). Crucially, the data are four times more likely under models which do not include the Grammaticality*TMS interaction (BF_excl_ = 4.058). Therefore, the additional analysis provide evidence against an effect of TMS over Broca’s area on the amplitude of the ESN component.

**Table 3:**
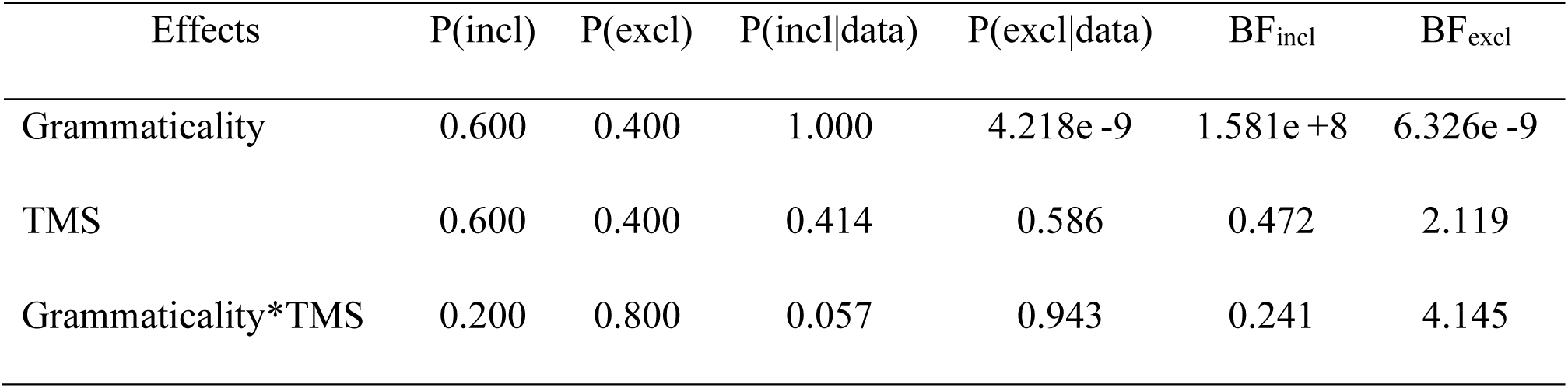
Summary of the analysis of the effects across all models. P(incl) = prior probability of including a predictor; P(excl) = prior probability of excluding a predictor; P(incl|data) = posterior probability of including a predictor; P(excl|data) = posterior probability of excluding a predictor; BF_incl_ = Bayes factor for including a predictor; BFexcl = Bayes factor for excluding a predictor.

### 3.4 ERP and induced electrical field simulation

The average intensity of the induced electrical fields in each ROI is summarized in Table 4. Within Broca’s area, the average electrical field was higher in BA45 (80.05 V/m) than BA44 (59.68 V/m). Within the SPL, the ROI in which TMS induced the highest electrical field were BA7PC (52.60 V/m), BA5L (41.66 V/m) and BA7A (41.33V/m). The reconstructed electrical fields for each subject, mapped to fsaverage space, are shown in Figures S9 and S10 in the Supplementary Materials.

**Table 4:** Mean and standard deviation of the induced electrical field (V/m) in the nine ROIs of interest

| Mean electrical field (SD) |  |  |  |
| --- | --- | --- | --- |
| Broca's area |  | Superior Parietal Lobe |  |
| BA44 | 59.68 (12.65) | BA5Ci | 15.76 (3.03) |
| BA45 | 80.05 (16.10) | BA5L | 41.66 (6.69) |
|  |  | BA5M | 17.52 (2.97) |
|  |  | BA7A | 41.33 (7.16) |
|  |  | BA7M | 13.02 (2.44) |
|  |  | BA7P | 23.29 (3.90) |
|  |  | BA7PC | 52.60 (12.15) |

Considering the Full ESN time-window, no significant correlation was found between Full ESN_BA44_ effect and the induced electrical field in BA44 (Table 5 and Figure 5, *r =* 0.142, *p* > 0.1, BF_01_ = 3.302, with median posterior δ = 0.128, 95% Credible Interval CI = [-0.239, 0.473]). The BF_01_ indicates that the data are 3.302 times more likely under the null hypothesis compared to the alternative one. Similarly, no significant correlation was found between Full ESN_BA44_ effect and the induced electrical field in BA45 (*r =* 0.114, *p* > 0.5, BF_01_ = 3.588, with median posterior δ = 0.103, 95% CI = [-0.264, 0.452]). No significant correlation was found between Full ESN_SPL_ effect and the induced electrical field in BA7PC (*r* = 0.011, *p* > 0.5, BF_01_ = 4.178, with median posterior δ = 0.010, 95% CI = [-0.352, 0.370]), in BA5L (*r* = 0.261, *p* > 0.1, BF_01_ = 1.845, with median posterior δ = 0.236, 95% CI = [-0.128, 0.560]) or in BA7A (*r* = 0.157, *p* > 0.1, BF_01_ = 3.126, with median posterior δ = 0.142, 95% CI = [-0.225, 0.484]). No significant correlation was found between Full ESN_SPL_ effect and the other SPL ROIs (see Table S1 in Supplementary Materials).

**Figure 5:**
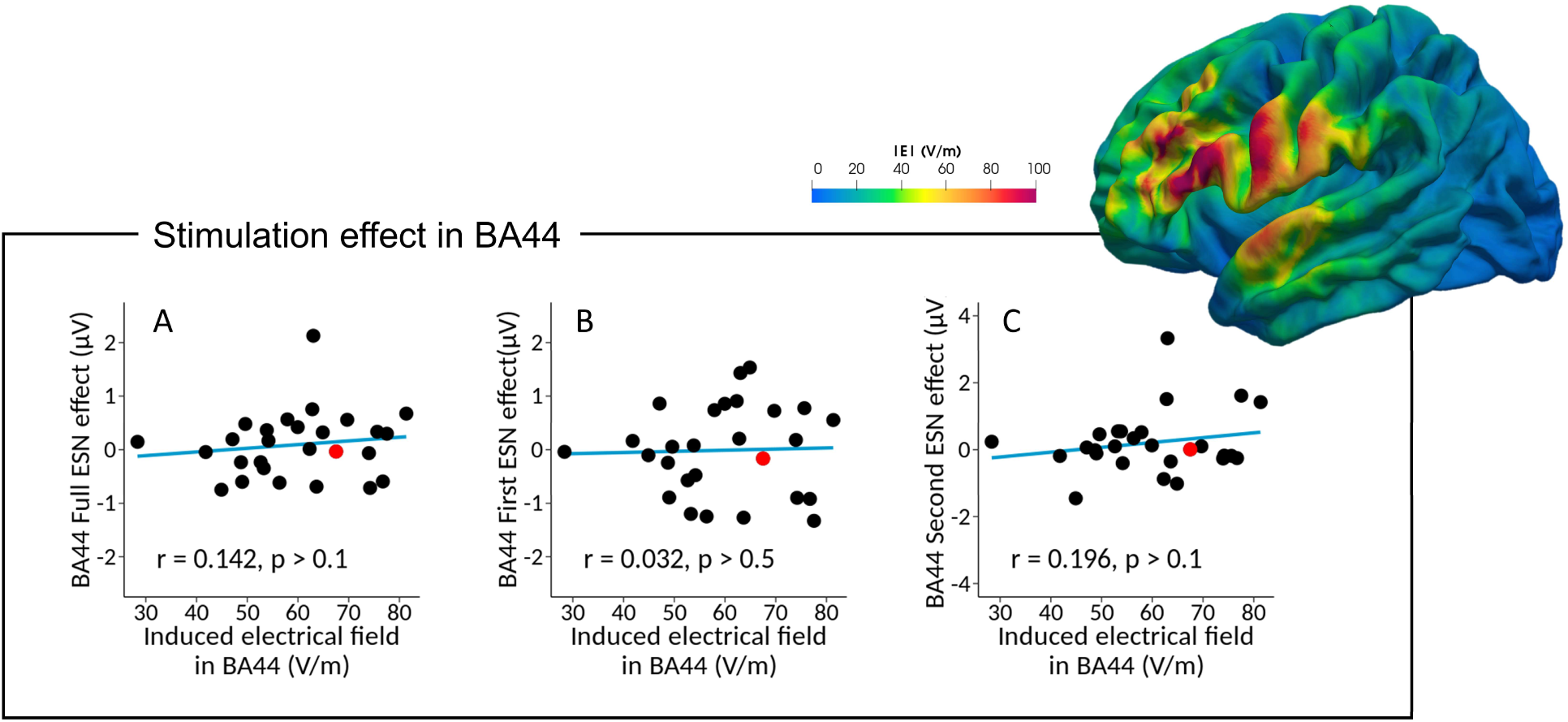
Correlation analysis between the Full ESN_BA44_ effect (Full ESN_BA44_ - Full ESN_sham_) and the induced electrical field in BA44 (A), together with separate correlation analyses for the First ESN_BA44_ effect (B) and Second ESN_BA44_ effect (C), respectively. The plotted brain illustrates the reconstructed TMS-induced electrical field from the BA44 session for a single subject, highlighted in red in the scatter plots.

**Table 5:** Correlational analysis between the induced electrical field in the subregions of Broca’s area and the three ESN effects of interest

| ESN effect | ROI eField | $r$ | $p$ | $BF_{10}$ | $BF_{01}$ |
| --- | --- | --- | --- | --- | --- |
| Full ESN<sub>BA44</sub> effect | BA44 | 0.142 | 0.480 | 0.303 | 3.302 |
| Full ESN<sub>BA44</sub> effect | BA45 | 0.114 | 0.570 | 0.279 | 3.588 |
| First ESN<sub>BA44</sub> effect | BA44 | 0.032 | 0.874 | 0.242 | 4.134 |
| First ESN<sub>BA44</sub> effect | BA45 | -0.006 | 0.975 | 0.239 | 4.182 |
| Second ESN<sub>BA44</sub> effect | BA44 | 0.196 | 0.327 | 0.378 | 2.648 |
| Second ESN<sub>BA44</sub> effect | BA45 | 0.120 | 0.549 | 0.283 | 3.529 |

Considering the first part of the ESN effect, no significant correlation was found between First ESN_BA44_ effect and the induced electrical field in BA44 (*r* = 0.032, *p* > 0.5, BF_01_ = 4.134, with median posterior δ = 0.029, 95% CI = [-0.334, 0.387]) and BA45 (*r* = -0.006, *p* > 0.5, BF_01_ = 4.182,with median posterior δ = -0.006, 95% CI = [-0.366, 0.356]). Similarly, no significant correlation was found between First ESN_SPL_ effect and the electrical field induced in BA7PC (*r* = -0.008, *p* > 0.5, BF_01_ = 4.182, with median posterior δ = -0.007, 95% CI = [-0.367, 0.355]), in BA5L (*r =* 0.294, *p* > 0.1, BF_01_ = 1.459, with median posterior δ = 0.267, 95% CI = [-0.095, 0.583]) or in BA7A (*r* = 0.149, *p* > 0.1, BF_01_ = 3.217, with median posterior δ = 0.135, 95% CI = [-0.233, 0.478]). No significant correlation was found between First ESN_SPL_ effect and the other SPL ROIs (see Table S2 in Supplementary Materials).

Finally, considering the second half of the ESN effect, no significant correlation was found between Second ESN_BA44_ effect and the induced electrical field in BA44 (*r* = 0.196, *p* > 0.1, BF_01_ = 2.648, with median posterior δ = 0.177, 95% CI = [-0.189, 0.513]) or in BA45 (*r* = 0.120, *p* > 0.5, BF_01_ = 3.529, with median posterior δ = 0.109, 95% CI = [-0.258, 0.456]). Similarly, no significant correlation was found between Second ESN_SPL_ effect and the electrical field induced in BA7PC (*r* = 0.014, *p* > 0.5, BF_01_ = 4.174, with median posterior δ = 0.013, 95% CI = [-0.349, 0.373]), in BA5L (*r =* 0.221, *p* > 0.1, BF_01_ = 2.337, with median posterior δ = 0.200, 95% CI = [-0.166, 0.531]) or in BA7A (*r* = 0.172, *p* > 0.1, BF_01_ = 2.949, with median posterior δ = 0.155, 95% CI = [-0.212, 0.495]). No significant correlation was found between Second ESN_SPL_ effect and the other SPL ROIs (see Table S3 in Supplementary Materials).

To summarize the analysis, even when accounting for the induced electrical field and the spatio- temporal profile of our ERP effect, our data show that TMS over Broca’s area did not affect the amplitude of the ESN component when inducing a virtual lesion during the online processing of the first word in our two-word paradigm.

## 4. DISCUSSION

Lesion studies provide evidence for a causal role of Broca’s area in syntactic composition, (Friederici et al., 1998, 1999), but leave open the question of whether this region is involved in predicting words’ grammatical categories or integrating them into constituents. A disruption of either of these processes would result in the absence of the ELAN in patients with left IFG lesion. In this study, we used TMS in healthy individuals to disentangle these processes by perturbating activity in Broca’s area during the first word (determiner or pronoun) of two-word phrasal/sentential structures. The high temporal resolution of the perturbation in our experiment allowed for the first time to specifically test the causal involvement of Broca’s area at the stage of syntactic prediction (determiner *→* prediction for a noun, pronoun *→* prediction for a verb). State-of-the-art modelling of the induced electrical field (Kuhnke et al., 2020; Weise et al., 2020) further quantified the impact of TMS in Broca’s area. Crucially, the online combination of EEG and TMS allowed to isolate the stimulation effect on both early automatic (ESN) and late controlled (late positivity) syntactic processes, further specifying the neuro-cognitive architecture of phrasal building. The present EEG-TMS data provided two main results. First, a main effect of Grammaticality revealed early automatic (ESN) and late controlled (late positivity) syntactic effects at the two-word level, functionally equivalent to those observed for longer and more complex stimuli (ELAN and P600, Friederici, 2002, 2011; Friederici & Kotz, 2003). Secondly, the absence of the critical Grammaticality*TMS interaction indicated that the transient disruption of Broca’s area at the stage of categorical prediction did not affect the generation of the ESN (prediction error, according to a predictive coding perspective). In the following sections, we discuss both findings in light of the previous literature.

### 4.1 Early and late main effects of grammaticality

The analysis of the main effect of Grammaticality revealed the presence of the ESN (approximately between 190 and 430 ms), followed by a late positivity (approximately between 440 and 800 ms). The presence of the ESN replicates previous work (Hasting & Kotz, 2008) and provides further evidence for an early analysis of categorical information at the most fundamental two-word level. The onset latency of the ESN in our experiment (∼200 ms) was slightly delayed compared to the original ESN study (Hasting & Kotz, 2008), in which the ERP waveforms for grammatical and ungrammatical constructions diverged already between 100 and 200 ms. Crucially, in the original ESN study the grammatical category was marked by the presence or absence of a single phoneme after the DVP (e.g., “Kegel_[DVP]_Ø”, *cone*, “kegel_[DVP]_t”, *bowls*) for the majority of the items (34/50 pairs of nouns and verbs). In our experiment, the grammatical category was always marked by a full syllable (e.g., “Fech_[DVP]_ter”, *fencer*, and “fech_[DVP]_tet”, *fences*), which unfolds over a longer time interval compared to a single phoneme. As a consequence, the detection of the grammatical violation is shifted in time. Another difference between the previous and present study is the offset time. While in the original ESN paradigm the effect lasted until 300 ms, our ESN effect lasted approximately 140 ms longer. There are two possible explanations for this discrepancy. On the one hand, considering that a full syllable and not a single phoneme marks the category in our stimulus set, the delayed offset time could simply be a consequence of the shift in onset latency of the ESN. On the other hand, the extended duration of our main effect might reflect the concatenation of two processes, indexed by a first anterior negativity (ESN) and a second N400. This pattern has previously been reported for agreement violations (Barber & Carreiras, 2005; Hanna et al., 2014; Jakuszeit et al., 2013) and marked categorical (Hasting et al., 2007) when realized at the two-word level. The presence of an N400 in agreement violations, where the meaning of grammatical and ungrammatical constructions is extremely similar (e.g., “a boat/*boats”), suggests that this component might reflect a process at the two-word level which is not purely semantic. Our stimuli match agreement violations paradigms with respect to the presence of a suffix indicating whether a given construction is grammatical or not. Therefore, the second negativity (N400) in our dataset could reflect an additional process in which a given suffix is compared against an expected one, which can be used to detect grammaticality for categorical violations overtly marked.

While functionally equivalent to the ELAN, the ESN does not show a left-lateralized topography. The early effect in our dataset has a rather non-lateralized and distributed topography, similar to the ESN reported by Hasting and Kotz (2008). In this regard, an fMRI study employing an adapted version of the original paradigm showed that categorical violations at the two-word level engage the bilateral temporal cortices, in addition to BA44 in the left hemisphere (Herrmann et al., 2012). These results are compatible with the presence of dipoles in both temporal cortices in ESN paradigms, which would explain the non-lateralized topography of the component. Interestingly, several ERP studies using two-word stimuli did not show a left-lateralized topography for categorical or agreement violation effects (Barber & Carreiras, 2005; Hasting & Kotz, 2008; Jakuszeit et al., 2013). Our results converge with these studies, indicating that the left-lateralized topography observed for syntactic effects in long sentences (Friederici, 2011) could become more central at the two-word level.

Finally, the ESN was followed by a late positivity, approximately between 440 and 800 ms. This late positivity aligns well with the profile of the P600 (Osterhout et al., 1994; Osterhout & Holcomb, 1993), an ERP component indexing repairing and re-analysis processes (Friederici, 2011). At the two-word level, the presence of a late positivity has been reported for agreement (Barber & Carreiras, 2005; Hasting & Kotz, 2008) and categorical (Jakuszeit et al., 2013) violations. This component was observed when a grammaticality judgement task was performed (agreement violations, Barber & Carreiras, 2005 and Hasting & Kotz, 2008) or during passive listening (categorical violations, Jakuszeit et al., 2013). However, when participants were actively distracted, this component was absent (Hasting & Kotz, 2008). In our experiment, participants were actively engaged in a repairing process, as they were performing a grammatical judgement task. Thus, the late positivity reflects a late and non-automatic process, as the P600 observed for longer stimuli (Anjia Hahne & Friederici, 1999). The present data converge with earlier studies demonstrating that the late syntactic processes observed with longer sentential stimuli (Anjia Hahne & Friederici, 1999) can be observed already at the minimal two-word level (Barber & Carreiras, 2005; Hasting & Kotz, 2008; Jakuszeit et al., 2013). Overall, our findings suggest that the recursivity that characterizes syntactic composition (Everaert et al., 2015; Friederici et al., 2017) can be observed at the neurophysiological level, with functionally equivalent processes at the basis of building both minimal phrases and more complex structures.

### 4.2 No evidence for Broca’s area’s causal role in categorical prediction

We perturbed activity in Broca’s area at the stage of syntactic prediction, by delivering TMS at the onset of a function word (“Ein”, *a*, and “Er”, *he*). We initially expected that disruption of Broca’s area would interfere with the formation of a categorical prediction (determiner → noun, pronoun → verb), leading to the absence of the ESN effect elicited by an ungrammatical continuation of the utterance (*determiner + verb, *pronoun + noun). However, the non-significant interaction between Grammaticality and TMS in our data does not support a causal role of Broca’s area in categorical prediction. At a more fine-grained level, modelling of the induced electrical field showed no relation between the ESN amplitude change and individual interference in Broca’s area. The additional Bayesian analysis provided initial evidence for the absence of the critical interaction effect, with the null hypothesis explaining our data approximately three times better than the alternative one. These results are compatible with two interpretations: either other brain regions outside of the left IFG are involved in generating syntactic categorical predictions, or automatic phrasal building does not rely on top-down expectations. We will discuss the first interpretation here, while we concentrate more closely on the second interpretation in the concluding paragraph of the discussion section.

In visual attention reading paradigm Bonhage and colleagues (2015) showed that categorical predictions engaged additional regions outside Broca’s area, such as the bilateral superior temporal cortices and insulae, the left frontal operculum, the intraparietal sulcus, the right caudate and the anterior cingulate. Some of these activations might reflect increased attentional demands triggered by the employed jabberwocky stimuli, which were used to highlight brain regions predicting an abstract category but not a specific word. Yet, other studies speak against the localization of categorical predictions in these regions. Activity in the frontal operculum and the insula increases as a function of the number of words presented, irrespectively of whether they are predictive or not (Zaccarella & Friederici, 2015a, 2015b). Similarly, the inferior parietal lobe shows increased activity for lists relative to constituents (Pallier et al., 2011), even if clear categorical predictions can be formed only in the latter type of stimuli. In this regard, direct comparison of sentences against lists showed that, once semantic processes are carefully subtracted, increased activation is found in BA44 only (Goucha & Friederici, 2015). Finally, conflicting evidence exists regarding an involvement of the superior temporal lobe in categorical predictions. Structural effects in the anterior temporal lobe (ATL) have been reported in the literature (Brennan & Pylkkänen, 2012; Brennan et al., 2016), and activity in this region has been linked to left-corner predictive parsing analysis (Brennan & Pylkkänen, 2012). However, these studies used narratives and sentences with real words, therefore it is unclear whether they isolated syntactic or semantic processes. Indeed, studies employing jabberwocky stimuli did not find a consistent involvement of the ATL in processing abstract categorical information (Goucha & Friederici, 2015; Matchin et al., 2019; Pallier et al., 2011; Zaccarella & Friederici, 2015a), and recent data point towards a role of this region in semantic/conceptual processing (Pylkkänen, 2020). The posterior temporal lobe has been shown to support top-down predictive processes in the semantic domain (Gastaldon et al., 2020), but it is unclear if this region subserves the automatic generation of categorical predictions. Recent fMRI and MEG data showed that the posterior superior temporal gyrus (pSTG) rather supports active syntactic predictive processes, which are generated at the sentential but not at phrasal level (Matchin et al., 2017, 2019). Since Bonhage and colleagues (2015) used sentence stimuli and included a long delay (five seconds) at the predictive stage, it cannot be excluded that the reported activity in the pSTG reflected a controlled process in this study. Such an active process cannot account for the fast and automatic nature of syntactic composition, and indeed lesion of the posterior temporal lobe does not affect the ELAN response (Friederici et al., 1998). Thus, while we only tested the causal involvement of Broca’s area in categorical prediction, previous functional and lesion data do not provide strong evidence for the involvement of the abovementioned brain regions in this process.

### 4.3 The role of Broca’s area role in syntactic composition

While our results do not support a causal role of Broca’s area in categorical predictions, they appear to be compatible with the alternative hypothesis that this region might be involved in the bottom-up integration of words into syntactic structures. Accordingly, in our experiment the ESN was not affected by TMS over Broca’s area simply because the stimulation occurred before this region was involved in the compositional process, as no syntactic rule could be evaluated on an isolated function word.

A first line of evidence supporting bottom-up syntactic composition in Broca’s area comes from studies which compared sentences and phrases against control conditions containing function words. Maguire and Frith (2004) showed increased activation in Broca’s area pars opercularis for sentences than lists, even if both conditions contained predictive function words. Similarly, Zaccarella and Friederici (2015) reported increased activity in BA44 not only for two-word pseudo- phrases relative to lists (e.g., “Diese Flirk”, *this flirk*, against “Apfel Flirk”, *apple flirk*), but also for determiner phrases compared to a single determiner (“Diese Flirk”, *this flirk,* against “Diese”, *this*). These studies support the notion that Broca’s area, and specifically BA44, is involved in the bottom-up integration of words into structures, as categorical predictions could be generated also in the control condition. Converging evidence comes from an fMRI study which investigated categorical violations at the two-word level (Herrmann et al., 2012). Herrmann and colleagues (2012) observed increased activity in BA44 for categorical violations (*pronoun + noun, *preposition + verb) compared to the grammatical items (pronoun + verb, preposition + noun). Crucially, the grammatical and ungrammatical items differed only in whether the second word violated a syntactic rule or not, as syntactic predictions triggered by the first word would in principle be present in both conditions. Therefore, the increased activation of BA44 in this experiment might reflect the bottom-up detection of an error, indexing that no grammatical rule to be applied is found. In this regard, recent electrocorticography and fMRI studies showed that the number of operations applied to integrate a word in the syntactic structure (bottom-up node count) correlated with activity in Broca’s area during language comprehension (Bhattasali et al., 2019; Nelson et al., 2017). Finally, preliminary comparison of bottom-up and predictive left-corner parsers showed that the former model provided a better account of activity in Broca’s area (Nelson et al., 2017).

In light of the discussion above, we advance the hypothesis that Broca’s area processes grammatical rules in a bottom-up fashion as words are incrementally encountered. Accordingly, words are temporarily stored until a grammatical rule can be applied to combine them into a syntactic structure. This notion is in line with the existence of distinct circuits maintaining words in memory and binding them into hierarchical structures (Iwabuchi et al., 2019; Makuuchi et al., 2009). At the two-word level, such a dissociation might be reflected in the distinct functional profile of the frontal operculum/adjacent insula and BA44 (Zaccarella & Friederici, 2015a, 2015b). The frontal operculum and insula increase their activity as a function of the number of words presented (Zaccarella & Friederici, 2015a, 2015b), while increased activation in BA44 is observed only when a grammatical rule can be invoked to combine two elements into a constituent (Zaccarella & Friederici, 2015a).

To the best of our knowledge, no study has directly tested the causal role of Broca’s area in bottom-up syntactic composition. However, evidence from agreement paradigms suggests that the left IFG might be involved in the bottom-up application of syntactic rules. Increased activity has been observed in the left IFG for agreement violations (Carreiras et al., 2010; Heim et al., 2010) and lesions in this region result in the absence of the ESN in this domain (Jakuszeit et al., 2013). While some authors linked the damage of the left IFG to the generation of predictions (Jakuszeit et al., 2013), the ESN would be absent even if the lesion affected the bottom-up evaluation of grammaticality. Data from a TMS study with two-word agreement violations support the latter hypothesis (Carreiras et al., 2012): stimulation of Broca’s area during the second word of a phrase causally affected bottom-up morphosyntactic processes, as indicated by a reduced advantage for a grammatical condition compared to an ungrammatical one.

Before concluding, we would like to point out that we are not suggesting that structural predictions are never generated, but rather that they do not necessarily constitute an automatic mechanism which incrementally builds syntactic structures (Matchin et al., 2017, 2019). In this respect, it is noteworthy considering whether predictive coding (Friston, 2003, 2005; Friston & Kiebel, 2009; Rao & Ballard, 1999) provides an adequate framework for syntactic composition. Grammar consists of a set of rules which are not probabilistic but deterministic – either something is correct or not – and which are not defined by the individuals. In other words, grammatical rules constitute a model which is not internal and is not constantly updated, contrary to the processes well described under a predictive coding perspective (Zaccarella et al., 2021). Therefore, automatic syntactic processes treat common and uncommon constructions equally, as long as they are grammatical (Friederici et al., 1996; Herrmann et al., 2009; Pulvermüller & Assadollahi, 2007). Individuals can construct internal probabilistic models of how likely it is that specific syntactic structures will be produced (Kroczek & Gunter, 2017), but the pure application of grammatical rules might just be a binary process: either something is correct or it is not. A similar dissociation seems to exist at the neural level, where brain regions sensitive to the probabilistic structural expectation appear to be located outside of the left-lateralized language network (Kroczek & Gunter, 2020).

## 5. LIMITATIONS

In line with the reviewed literature, we have proposed a bottom-up integration role for this region in syntactic composition. An alternative account may posit that the TMS effect in the region might have been either short-lived or rapidly compensated, thus only minimally affecting the prediction phase in our study. Evidence from online (Klaus & Hartwigsen, 2019; Kuhnke et al., 2017; Meyer et al., 2018) and offline (Acheson & Hagoort, 2013) TMS studies, however, seems to speak against this explanation, since TMS application over Broca’s area has been conversely shown to affect linguistic processing. The estimation of the TMS-induced electrical fields in our participants is a first-time quantification of the realized IFG stimulation. To the best of our knowledge, this new perturbation quantification (Weise et al., 2020) has not been used elsewhere in the TMS literature on the left IFG. However, a recent study with stimulation of the parietal lobe showed that induced electrical fields weaker than the ones reported in our study were sufficient to induce a TMS effect on cognitive processes (Kuhnke et al., 2020). Thus, the available evidence suggests that the stimulation protocol and intensities used in the present study may have been well-suited for disrupting Broca’s area functioning at the predictive phase. Future studies targeting syntactic processes in Broca’s area, possibly exploiting novel methods for estimating TMS effects on neural processing (Kuhnke et al., 2020; Numssen et al., 2021; Weise et al., 2020), might provide useful insights, either by replicating the present results or by providing evidence for alternative hypotheses.

## 6. CONCLUSIONS

In this EEG-TMS study, we tested whether Broca’s area is causally involved during the categorical prediction phase in two-word phrasal/sentential constructions. Contrary to our hypothesis, perturbation of Broca’s area at the predictive stage did not affect the ERP correlates of basic syntactic composition. Our data are compatible with the proposal that phrasal building in Broca’s area might be accomplished in a bottom-up fashion (Bhattasali et al., 2019; Nelson et al., 2017), with words being integrated into constituents whilst the linguistic stream unfolds. They also converge on fMRI and TMS data implying bottom-up evaluation of grammatical rules in this region (Carreiras et al., 2010, 2012; Heim et al., 2010; Herrmann et al., 2012; Zaccarella & Friederici, 2015a). As such, future studies addressing this neurocognitive hypothesis are awaited to provide further insights into the mechanisms of incremental linguistic composition.

## ACKNOWLEDGMENTS

We wish to thank Philipp Kuhnke for assisting in the calculation of the target coordinates in subject space, Ina Koch and Joëlle Schroën for their help during data acquisition, Maren Grigutsch and Burkhard Maess for the fruitful discussions on the pre-processing pipeline, Giorgio Papitto, Elena Pyatigorskaya and Patrick Trettenbrein for their insightful comments on a previous version of this manuscript.

## CREDIT AUTHOR STATEMENT

**Matteo Maran:** Conceptualization, Methodology, Formal analysis, Investigation, Data curation, Writing - Original Draft, Writing - Review & Editing, Visualization; **Ole Numssen:** Formal Analysis (Electrical field simulations), Writing - Review & Editing, Visualization (Electrical field simulations); **Gesa Hartwigsen:** Resources, Writing - Review & Editing; **Angela D. Friederici**: Conceptualization, Resources, Writing - Review & Editing, Funding acquisition, Supervision; **Emiliano Zaccarella**: Conceptualization, Writing - Review & Editing, Visualization, Supervision.

## Footnotes

1 The symbol * is conventionally used in theoretical linguistics to indicate an ungrammatical construction.

2 Ø denotes a so-called zero form, i.e., an absence of a suffix.

3 In the case of the interaction, the independent variable to be considered is the grammaticality effect within each TMS condition, which has three levels: BA44 grammaticality effect, SPL grammaticality effect, sham grammaticality effect.

4 The coordinates used as target for stimulating BA44 (Zaccarella & Friederici, 2015a) lie indeed in the most anterior and ventral part of the region, very close to BA45.

5 We describe the temporal and spatial extent only in approximate terms, as recommended by methodological papers on this topic (Maris, 2012; Maris & Oostenveld, 2007; Sassenhagen & Draschkow, 2019).

