## Supplementary Materials for "Towards a causal role of Broca’s area in language: A TMS-EEG study on syntactic prediction"

[b]International Max Planck Research School on Neuroscience of Communication: Function, Structure, and Plasticity, Stephanstraße 1a, 04103, Leipzig, Germany

[c]Max Planck Institute for Human Cognitive and Brain Sciences, Lise Meitner Research Group Cognition and Plasticity, Stephanstraße 1a, 04103, Leipzig, Germany

#### SUPPLEMENTARY MATERIALS

##### Response cue colours

Relative luminance of the red and green colours used as response cues was calculated implementing the formula defined in the Web Content Accessibility Guidelines (WCAG) 2.0 (https://www.w3.org/TR/WCAG20/Overview.html#sRGB):

$L = 0.2126*R + 0.7152*G + 0.0722*B$

R, G and B are calculated as follows:

1. RsRGB = R8bit/255; If RsRGB is smaller or equal to 0.03928, R = RsRGB/12.92, otherwise R = ((RsRGB+0.055)/1.055) ^ 2.4;
2. GsRGB = G8bit/255; If GsRGB is smaller or equal to 0.03928, G = GsRGB/12.92, otherwise G = ((GsRGB+0.055)/1.055) ^ 2.4;
3. BsRGB = B8bit/255; If BsRGB is smaller or equal to 0.03928, B = BsRGB/12.92, otherwise B = ((BsRGB+0.055)/1.055) ^ 2.4.

### SUPPLEMENTARY MATERIALS - FIGURES

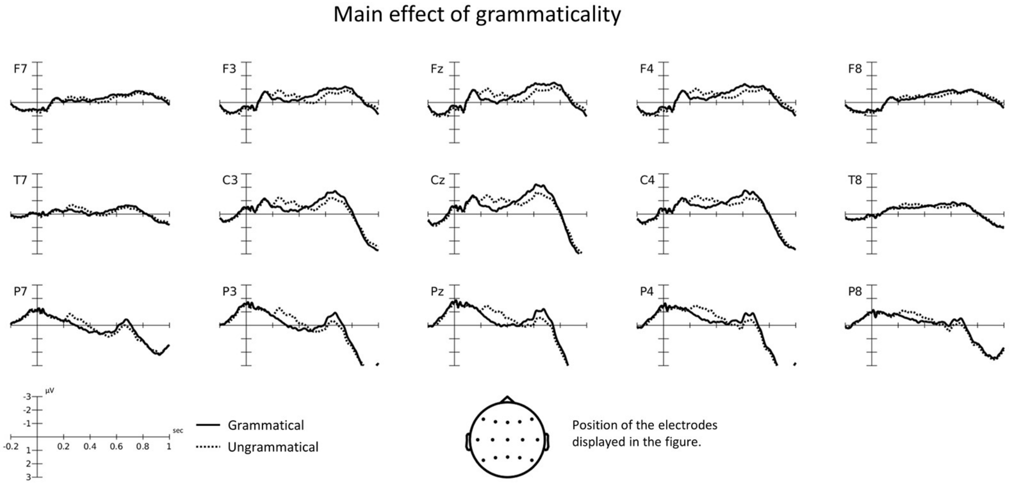


***Figure S1:*** *Figure S1 displays the ERP waveforms for the main effect of grammaticality at selected electrodes.*

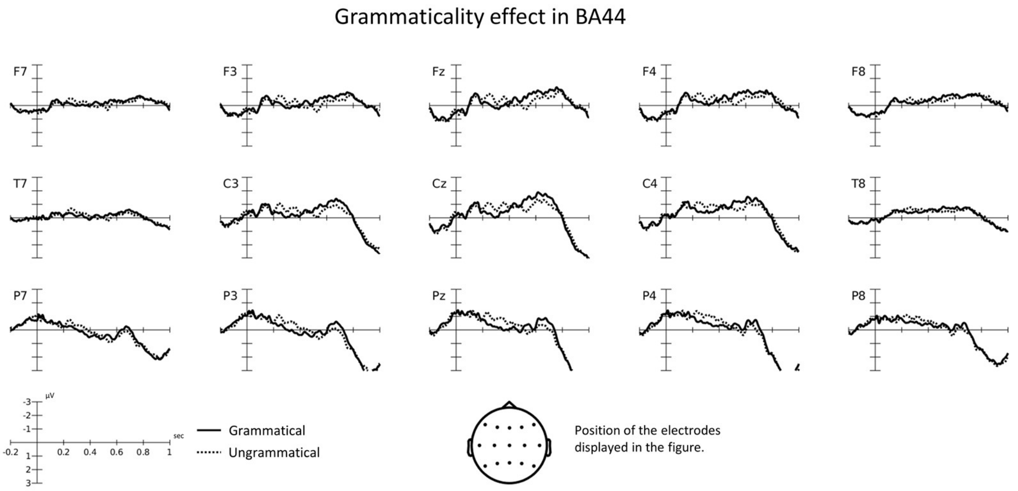


***Figure S2:*** *Figure S2 displays the ERP waveforms for the grammaticality effect for the TMS condition BA44 at selected electrodes.*

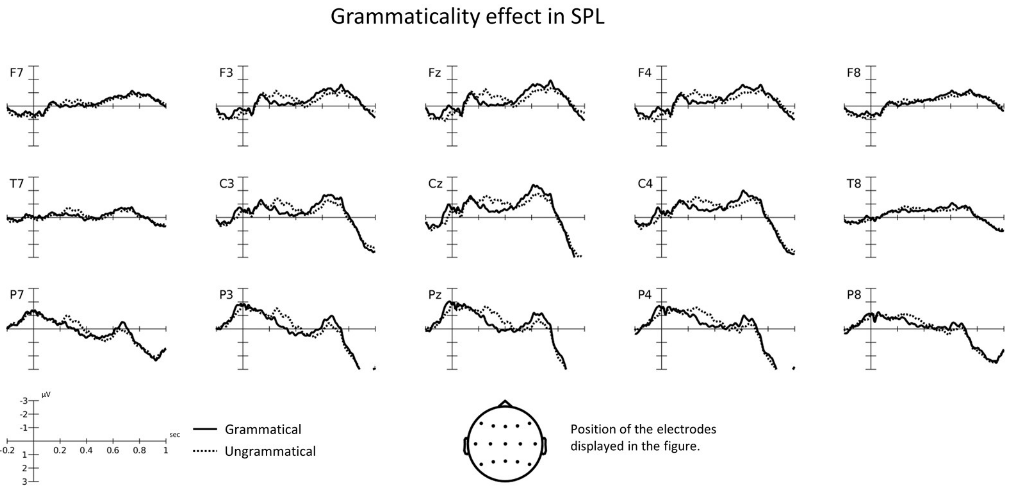


***Figure S3:*** *Figure S3 displays the ERP waveforms for the grammaticality effect for the TMS condition SPL at selected electrodes.*

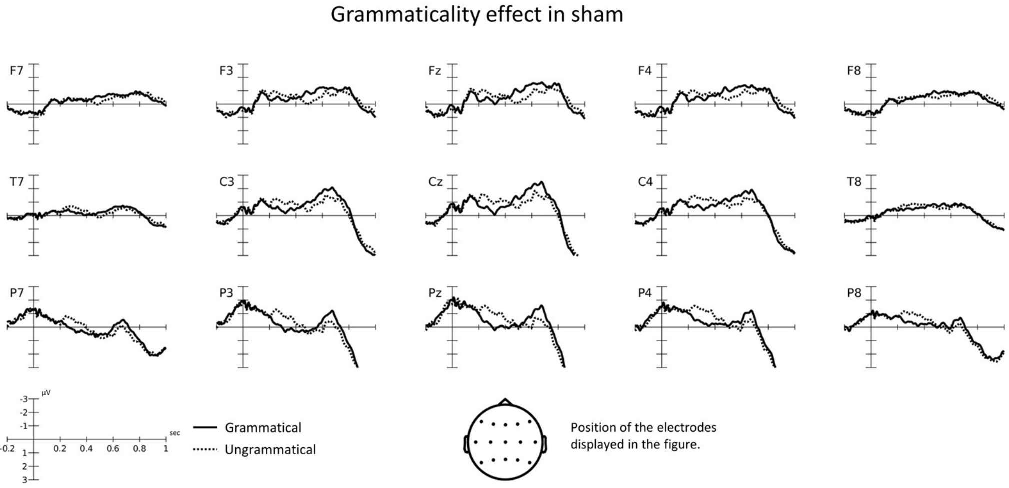


***Figure S4:*** *Figure S4 displays the ERP waveforms for the grammaticality effect for the TMS condition sham at selected electrodes.*

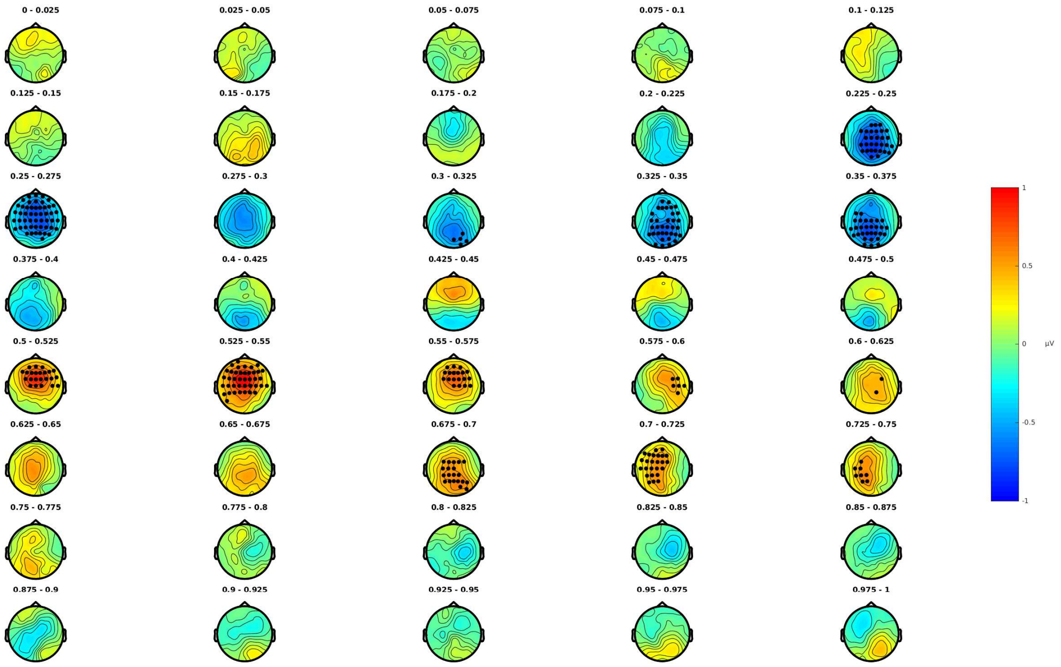


***Figure S5:*** *Figure S5 displays the topography of the Grammaticality effect for the BA44 condition. The electrodes and time-points mostly contributing to the significance of the negative and positive clusters are highlighted.*

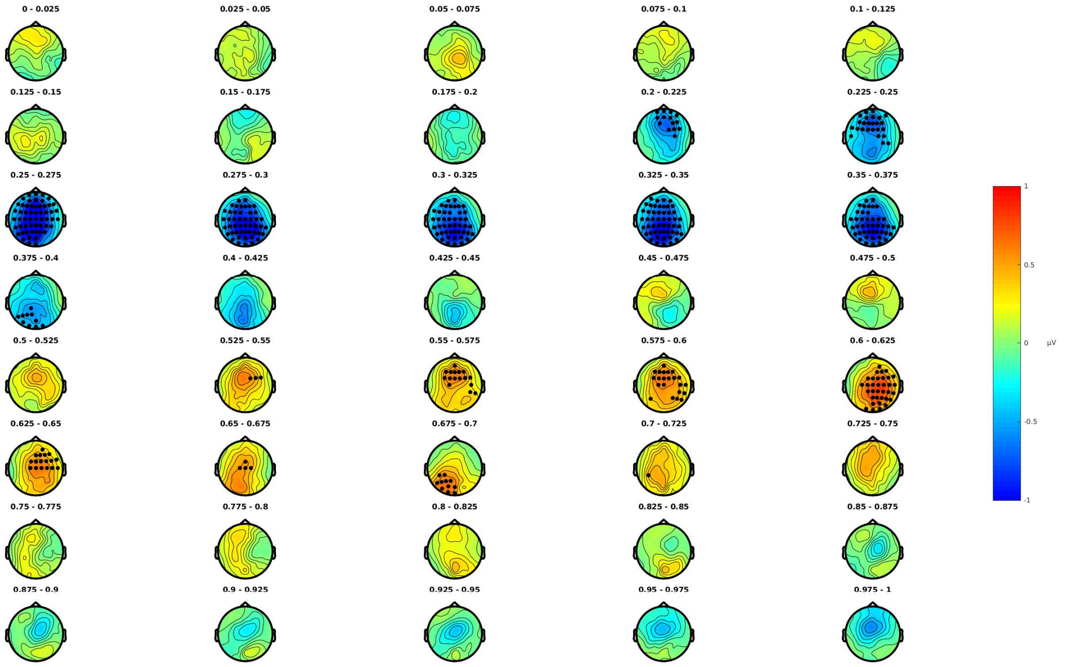


***Figure S6:*** *Figure S6 displays the topography of the grammaticality effect for the SPL condition. The electrodes and time-points mostly contributing to the significance of the negative and positive clusters are highlighted.*

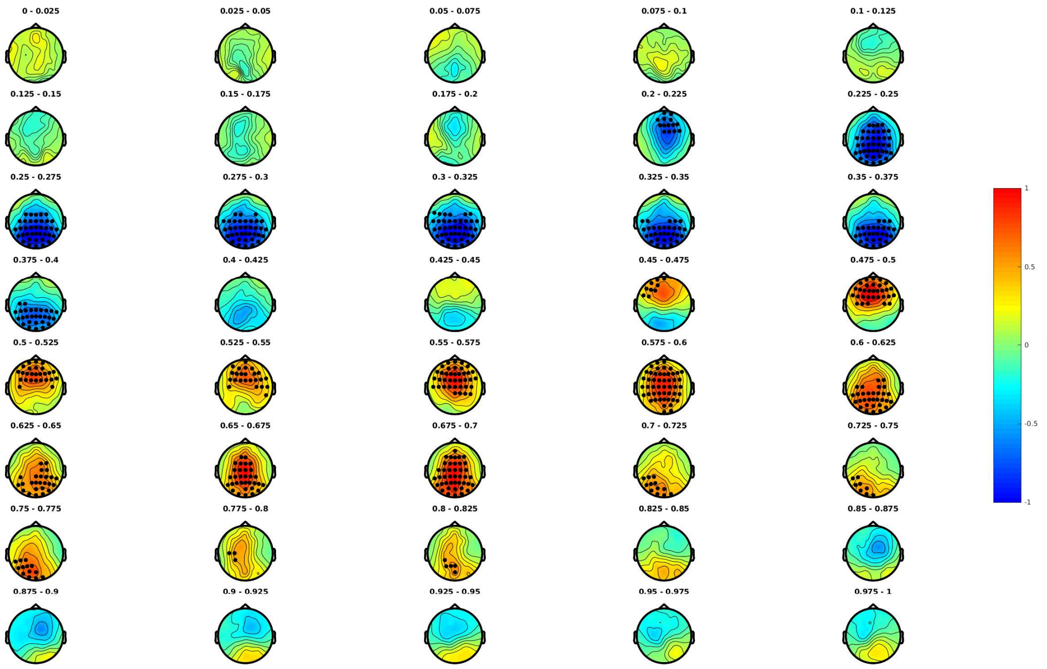


***Figure S7:*** *Figure S7 displays the topography of the grammaticality effect for the sham condition. The electrodes and time-points mostly contributing to the significance of the negative and positive clusters are highlighted.*

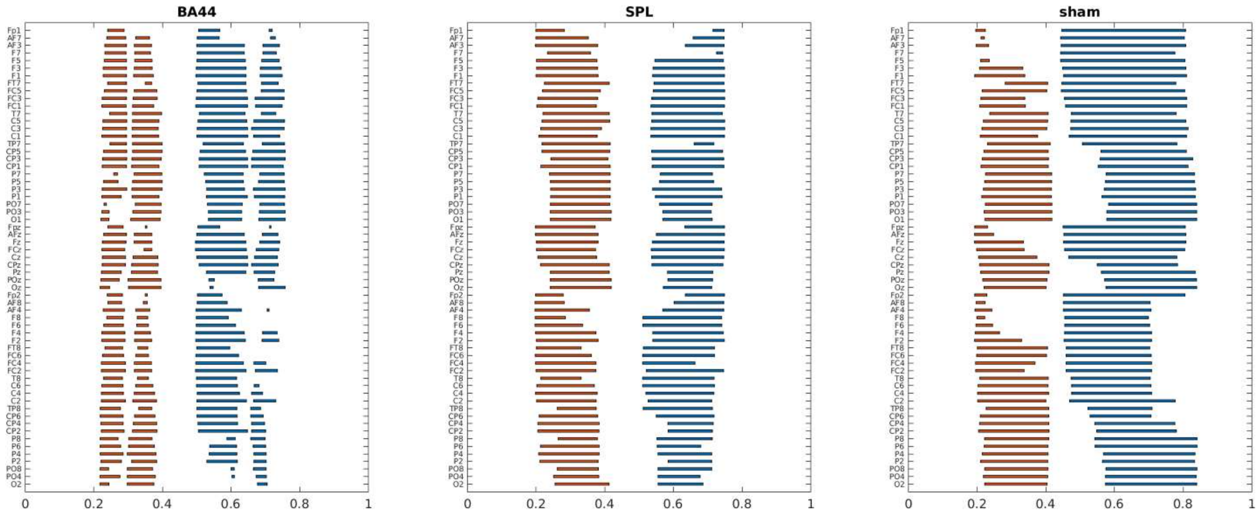


***Figure S8:*** *Figure S8 displays the spatial (electrode) and temporal (seconds) extent of the negative (red) and positive (blue) clusters for the Grammaticality effect in each of the TMS conditions.*

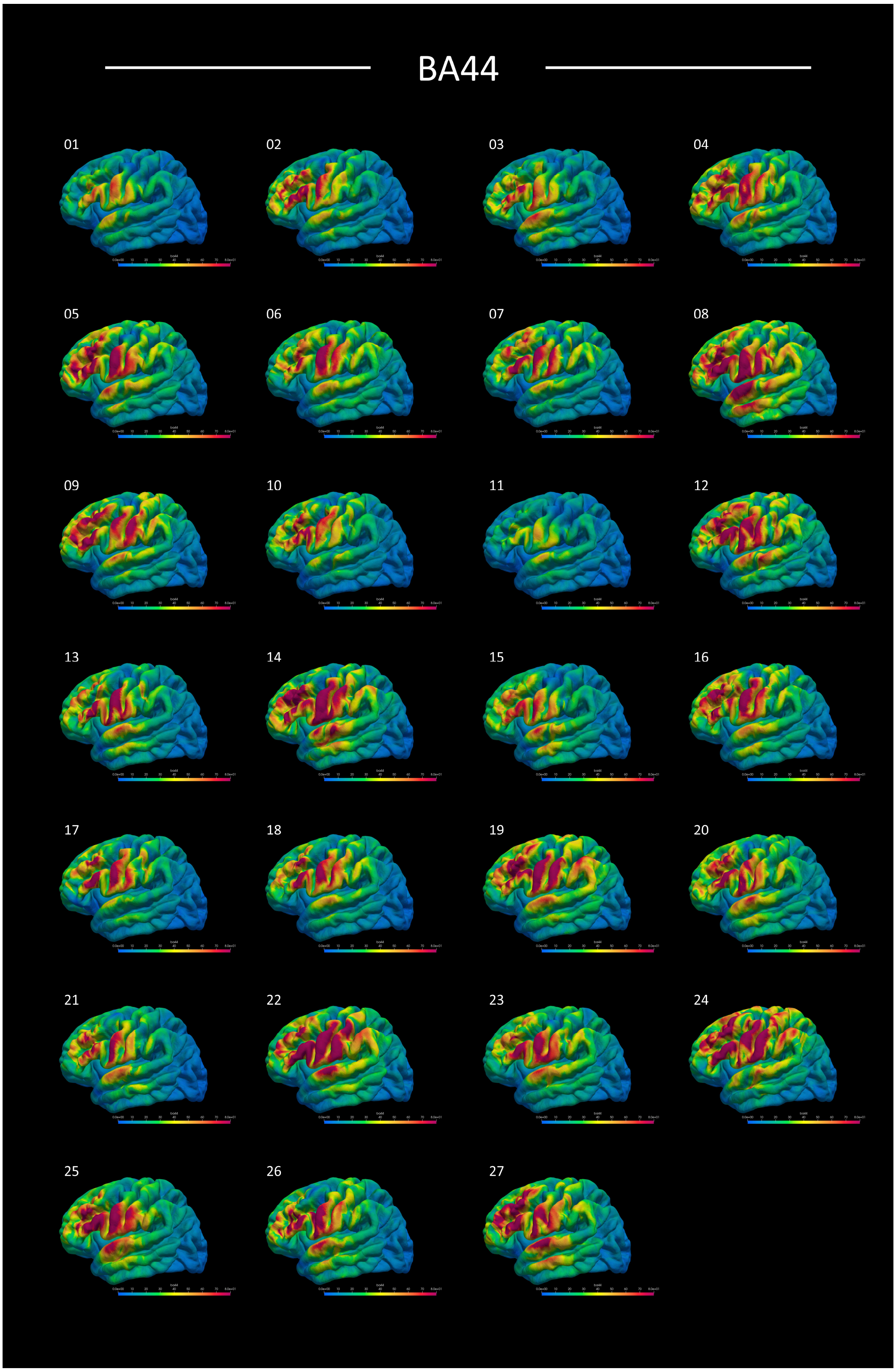


***Figure S9:*** *Visualization of the induced electric field magnitude for the BA44 TMS condition for each of the 27 subjects included in the analyses.*

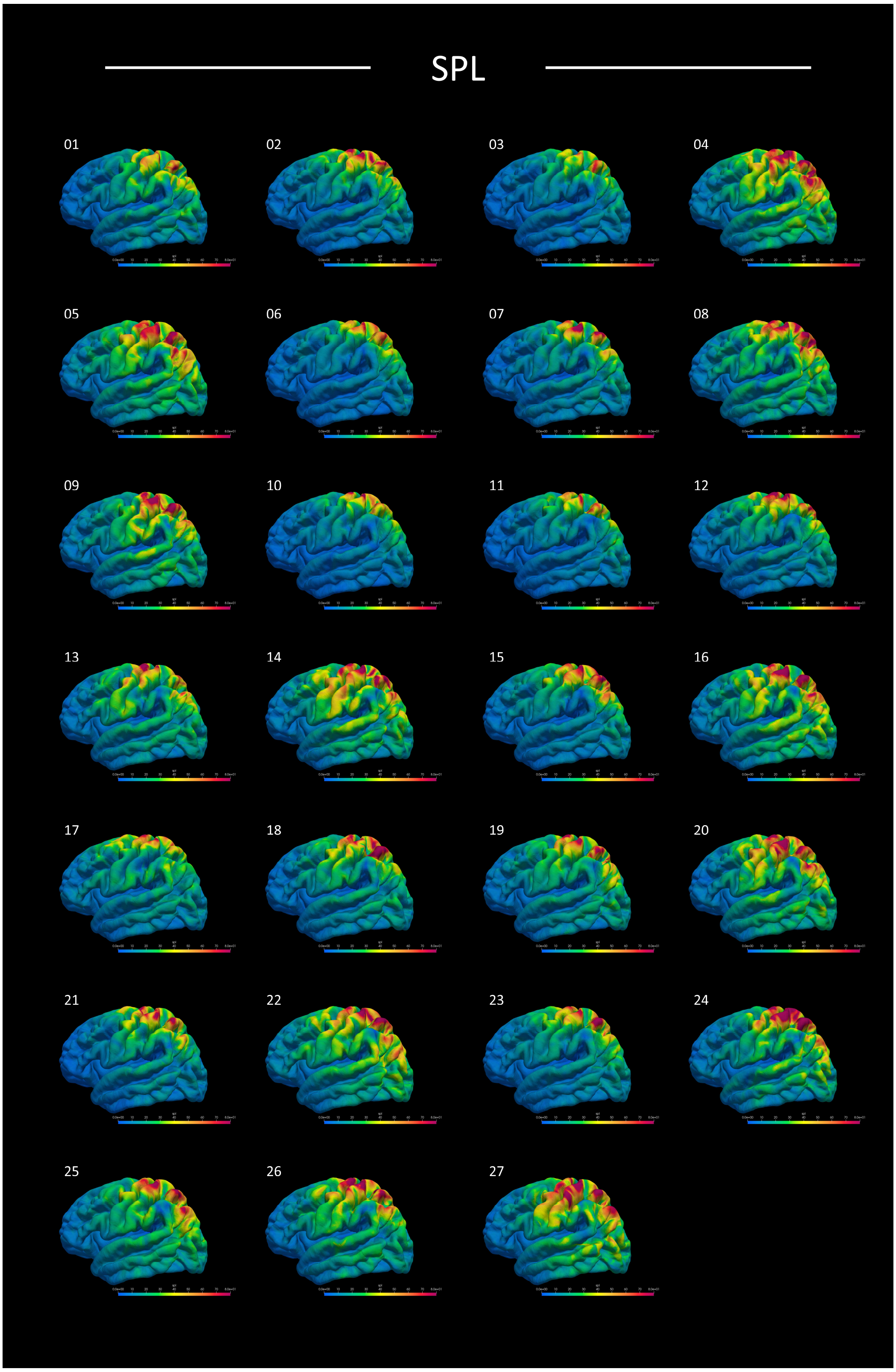


***Figure S10:*** *Visualization of the induced electric field magnitude for the SPL TMS condition for each of the 27 subjects included in the analyses.*

**SUPPLEMENTARY MATERIALS - TABLES**

| ESN effect | ROI eField | *r* | *p* | $BF_{10}$ | $BF_{01}$ |
|---|---|---|---|---|---|
| Full $ESN_{SPL}$ effect | BA5Ci | 0.095 | 0.639 | 0.265 | 3.769 |
| Full $ESN_{SPL}$ effect | BA5L | 0.261 | 0.189 | 0.542 | 1.845 |
| Full $ESN_{SPL}$ effect | BA5M | 0.150 | 0.454 | 0.312 | 3.206 |
| Full $ESN_{SPL}$ effect | BA7A | 0.157 | 0.434 | 0.320 | 3.126 |
| Full $ESN_{SPL}$ effect | BA7M | -0.021 | 0.916 | 0.240 | 4.163 |
| Full $ESN_{SPL}$ effect | BA7P | 0.089 | 0.658 | 0.262 | 3.812 |
| Full $ESN_{SPL}$ effect | BA7PC | 0.011 | 0.956 | 0.239 | 4.178 |

***Table S1:*** *Correlational analysis between the Full $ESN_{SPL}$ effect and the induced electrical field in the subregions of the SPL.*

| ESN effect | ROI eField | *r* | *p* | $BF_{10}$ | $BF_{01}$ |
|---|---|---|---|---|---|
| First $ESN_{SPL}$ effect | BA5Ci | 0.104 | 0.604 | 0.272 | 3.682 |
| First $ESN_{SPL}$ effect | BA5L | 0.294 | 0.137 | 0.685 | 1.459 |
| First $ESN_{SPL}$ effect | BA5M | 0.121 | 0.546 | 0.284 | 3.520 |
| First $ESN_{SPL}$ effect | BA7A | 0.149 | 0.457 | 0.311 | 3.217 |
| First $ESN_{SPL}$ effect | BA7M | -0.115 | 0.569 | 0.279 | 3.585 |
| First $ESN_{SPL}$ effect | BA7P | 0.079 | 0.696 | 0.257 | 3.891 |
| First $ESN_{SPL}$ effect | BA7PC | -0.008 | 0.970 | 0.239 | 4.182 |

***Table S2:*** *Correlational analysis between the First $ESN_{SPL}$ effect and the induced electrical field in the subregions of the SPL.*

| ESN effect | ROI eField | *r* | *p* | $BF_{10}$ | $BF_{01}$ |
|---|---|---|---|---|---|
| Second $ESN_{SPL}$ effect | BA5Ci | 0.124 | 0.538 | 0.286 | 3.493 |
| Second $ESN_{SPL}$ effect | BA5L | 0.221 | 0.268 | 0.428 | 2.337 |
| Second $ESN_{SPL}$ effect | BA5M | 0.218 | 0.275 | 0.421 | 2.377 |
| Second $ESN_{SPL}$ effect | BA7A | 0.172 | 0.391 | 0.339 | 2.949 |
| Second $ESN_{SPL}$ effect | BA7M | 0.119 | 0.555 | 0.282 | 3.546 |
| Second $ESN_{SPL}$ effect | BA7P | 0.096 | 0.634 | 0.266 | 3.757 |
| Second $ESN_{SPL}$ effect | BA7PC | 0.014 | 0.943 | 0.240 | 4.174 |

***Table S3:*** *Correlational analysis between the Second $ESN_{SPL}$ effect and the induced electrical field in the subregions of the SPL.*
